# TRANsCre-DIONE transdifferentiates scar-forming reactive astrocytes into functional motor neurons

**DOI:** 10.1101/2020.07.24.215160

**Authors:** Heeyoung An, Hye-Lan Lee, Doo-Wan Cho, Jinpyo Hong, Hye Yeong Lee, Jung Moo Lee, Junsung Woo, Jaekwang Lee, MinGu Park, Young-Su Yang, Su-Cheol Han, Yoon Ha, C. Justin Lee

## Abstract

In spinal cord injury (SCI), the scar-forming reactive astrocytes with upregulated GFAP proliferate aberrantly near the injury site, allowing themselves as a prime target for transdifferentiation into neurons to replenish dead neurons. However, the conventional use of *GFAP* promoter to target reactive astrocytes has two inherent problems: inadvertent conversion of normal astrocytes and low efficiency due to progressive weakening of promoter activity during transdifferentiation. Here, we report that the scar-forming reactive astrocytes are selectively transdifferentiated into neurons with 87% efficiency and 96% specificity via TRANsCre-DIONE, a combination of the split-Cre system under two different promoters of *GFAP* and *Lcn2* and a Cre-loxP-dependent inversion and expression of *Neurog2* under the strong *EF1α* promoter. After SCI, TRANsCre-DIONE caused transdifferentiation into Isl1-positive motor neurons, reduced astrogliosis, enhanced regeneration in surrounding cells, and a significant motor recovery. Our study proposes TRANsCre-DIONE as the next-generation therapeutic approach for patients suffering from SCI.

**Highlights:** TRANsCre-DIONE converts reactive astrocyte into neuron by over-expression of *Neurog2* Reactive astrocytes are targeted using split-Cre under two promoters, *GFAP* and *Lcn2* TRANsCre-DIONE reduces reactivity, replaces dead neurons and alleviates symptom of SCI Transdifferentiated-neurons are GABA+ in the striatum and Isl1+ in the spinal cord

## INTRODUCTION

Spinal cord injury (SCI) induces irreversible neuronal death and glial scar formation, which results in permanent motor and sensory dysfunction (Ahuja et al., 2017; Cregg et al., 2014; Su et al., 2014). Although many attempts have been set forth to regenerate damaged spinal cord, currently doctors can prescribe medications only to relieve pain to SCI patients. There is no drug or therapeutic tool to help regenerate neurons and cause functional recovery, other than strenuous rehabilitation regimes (Alizadeh et al., 2019; Assinck et al., 2017; Nagoshi and Okano, 2018). The much-hoped-for stem cell therapy still has a long way to go in clinics due to its issues with low efficacy and safety. The safety issues include tumor formation, immune rejection and unnecessary pain by reprogrammed neurons forming improper neural connections (Abad et al., 2013; Yin et al., 2019). These serious risks are majorly caused by transplantation of induced stem cells which still might have pluripotency (Li and Chen, 2016).

To overcome the limitations of transplantation, the concept of transdifferentiation or direct reprogramming of one type of differentiated resident cells into other cell types has been proposed (Li and Chen, 2016; Xu et al., 2015). It has been reported that diverse cell types, including fibroblast, microglia, and astrocyte can be transdifferentiated into neurons using various transcription factors (Guo et al., 2014; Heinrich et al., 2010; Matsuda et al., 2019; Takahashi et al., 2007; Victor et al., 2014). Among these cell types, the glial fibrillary acidic protein (GFAP)-positive astrocytes are known as the most abundant cell type in the brain (Barres, 1991). Under normal conditions, astrocytes provide various neurotrophic factors for neuronal growth and survival, regulate ionic homeostasis by taking up potassium ions and glutamate, and engage in synaptic transmission and plasticity by releasing gliotransmitters (De Pitta et al., 2011). However, under pathological conditions such as in SCI, these normal astrocytes transform at the site of injury into reactive astrocytes and become hypertrophied and proliferative, especially under a severe condition (Chun and Lee, 2018), to eventually form a glial scar (Anderson et al., 2016; Burda and Sofroniew, 2014; Okada et al., 2018). The severely reactive astrocytes over-express oxidizing enzymes such as MAO-B to cause oxidative stress and become highly toxic to neighboring neurons (Chun et al., 2018). Therefore, these proliferating and scar-forming severely reactive astrocytes can be a prime target for transdifferentiation into neurons to remove these toxic severely reactive astrocytes and replace dead neurons after injury.

*GFAP* promoter has been widely used to both target astrocytes and drive expression of transcription factors for transdifferentiation into neurons (Brulet et al., 2017; Guo et al., 2014; Heinrich et al., 2010; Mattugini et al., 2019; Su et al., 2014). Despite the universal application of the *GFAP* promoter (Brenner et al., 1994), the efficiency of transdifferentiation *in vivo* has been shown to be lower than 30% in a mouse model of spinal cord injury (Su et al., 2014). The low efficiency is probably due to most of the astrocytes being prematurely terminated of transdifferentiation by the reduced expression of a transcription factor under *GFAP* promoter as the *GFAP* promoter activity is downregulated by the conversion of astrocytes into neurons. Therefore, the use of *GFAP* promoter to drive the expression of a transcription factor in astrocytes for transdifferentiation may not be optimal. More importantly, using solely *GFAP* promoter is not an ideal strategy for targeting reactive astrocytes because GFAP is also expressed in some neural progenitor cells as well as normal astrocytes (Liu et al., 2010; Park et al., 2018). This raises a possibility that some neural progenitor cells and normal astrocytes might unintentionally transdifferentiate into neuron-like cells. In fact, although a recent study reported a high transdifferentiation efficiency of 80% with only *GFAP* promoter to target reactive astrocytes in the brain of Alzheimer’s disease and Huntington’s disease mouse models (Guo et al., 2014; Wu et al., 2020), the apparent high efficiency might have been over-estimated by an undesirable targeting of neural progenitor cells and normal astrocytes. Therefore, to target only reactive astrocytes while sparing neural progenitor cells and normal astrocytes, it is essential to utilize one or more reactive-astrocyte-specific promoters such as *Lcn2* (Bi et al., 2013) and *iNOS* (Walsh et al., 2004; Zhao et al., 1998) in combination with *GFAP* promoter. Moreover, to continuously drive the expression of a transcription factor even in the transdifferentiated state, it is necessary to utilize a strong universal promoter, whose activity is maintained high regardless of the cell types. To prevent an inadvertent expression of a transcription factor in untargeted cells under a strong universal promoter, the gene of transcription factor should remain in an inactive form, whereas it should be switched to an active form at once in targeted cells. To address these requirements, we have utilized the split-Cre system (Hirrlinger et al., 2009a; Hirrlinger et al., 2009b) under two separate promoters for reactive astrocytes, which will switch on a gene of transcription factor by Cre/DIO (double floxed inversed orientation)-dependent inversion (Tronche et al., 2002). The inverted gene of transcription factor is then continuously expressed as it is driven by a strong universal promoter *EF1α* (Wang et al., 2017).

To transdifferentiate reactive astrocytes into motor neurons in spinal cord injury model, we have considered various transcription factors that have been used to transdifferentiate astrocytes to neurons. To transdifferentiate fibroblasts into neurons directly, achaete-scute family bHLH transcription factor 1 (A*SCL1)* has been included in combination with other transcription factors such as *Sox2, Oct4*, and *Dlx1* (Liu et al., 2016). To transdifferentiate astrocytes into neurons, *Neurog2* and its downstream factor, *NeuroD1* (Seo et al., 2007), have been sequentially introduced as a single transcription factor *in vitro* (Berninger et al., 2007; Heinrich et al., 2010) and *in vivo* (Guo et al., 2014; Hu et al., 2019). To transdifferentiate fibroblasts into the Isl1-positive motor neurons, *Neurog2* has been included in combinations with other transcription factors (Liu et al., 2016; Tang et al., 2017). However, *Neurog2* has not been used for transdifferentiation of reactive astrocytes into motor neurons in spinal cord injury model. On the other hand, *NeuroD1* has been used frequently for transdifferentiation of astrocytes into glutamatergic neurons in combination with other transcription factors (Wang et al., 2016), but rarely for transdifferentiation into motor neurons in combination with five other transcription factors (Wang et al., 2016). Only *Sox2* in combination with other factors has been used for transdifferentiation of astrocytes to neurons in spinal cord injury model (Wang et al., 2016). However, the study did not report any functional recovery after injury (Wang et al., 2016). Based on these previous findings, we selected *Neurog2* as an optimal single transcription factor for transdifferentiation of reactive astrocytes into motor neurons. In this study, we employ **TRANsCre-DIONE** (**T**ransdifferentiation of **R**eactive **A**strocytes to **N**eurons by **s**plit-**Cre** using GFAP and Lcn2 or iNOS turning on of **DIO**-**N**eurogenin2 under **E**F1a promoter), a unique combination of split-Cre under two separate promoters for reactive astrocytes and Cre/DIO-dependent inversion of *Neurog2* under a strong universal promoter of *EF1α* to transdifferentiate scar-forming reactive astrocytes into Isl1-positive motor neurons in hope for functional recovery in mouse model of spinal cord injury.

## RESULT

### The genetic strategy of transdifferentiation in the brain of a mouse and non-human primate

The scar-forming reactive astrocytes exhibit upregulation of *GFAP* and neuroinflammation-related genes such as lipocalin2 (*Lcn2*) and induced nitric oxide synthase (*iNOS*) (Walsh et al., 2004; Zhao et al., 1998). To express the transcription factor in reactive astrocytes more specifically, we introduced split-Cre which is the split-form of the whole improved Cre (iCre), divided into two parts composed of N-terminal part of the iCre (Ncre) and C-terminal part of the iCre (Ccre) (Hirrlinger et al., 2009a; Hirrlinger et al., 2009b), under two different promoters. To confirm the induction of the gene products of *Lcn2* and *iNOS* in the reactive astrocytes, we immunostained the injured tissues with antibodies against Lcn2 and iNOS. To apply the traumatic injury to the brain of the mouse and monkey, the 32-gauge and 23-gauge needle was used for virus injection, respectively (Figure 1A). To induce an injury to the mouse spinal cord, a compression injury was introduced by self-closing forceps for 3 s followed by virus injections into the two points next to the injury site (Figure 1A). The GFAP signal increased in both mouse and monkey when the tissues were injured. Lcn2 was induced in the injured mouse brain and spinal cord (Figure 1B) while only the iNOS signal was induced without the Lcn2 signal in the injured monkey’s brain tissues (Figure 1B). Lcn2 and iNOS were never observed in the uninjured tissues (Figure S1). Along with these results, for gene expression in reactive astrocytes specifically, *Lcn2* was used with *GFAP* as promoters of split-Cre in the mouse brain and spinal cord, and *iNOS* and *GFAP* were used for the promoters of split-Cre in the monkey’s brain (Figure 1C). All genes were cloned and packaged into Adeno-associated virus (AAV) for broad and robust gene expression in the brain and spinal cord. *Neurog2* was expressed only in reactive astrocytes because both sides of the split-Cre can be expressed only in reactive astrocytes to make a functional Cre. Thus, we used TRANsCre-DIONE, composed of pAAV-EF1α::DIO-Neurog2-IRES-GFP (DIO-Neurog2) with pAAV-Lcn2::NCre and pAAV-GFAP::CCre in the mouse and pAAV-iNOS::CCre and pAAV-GFAP::NCre in the monkey brain (Figure 1C). In addition, GFP fluorescence (under *IRES*; Internal Ribosome Entry Site) was used as a reporter for easy monitoring of TRANsCre-DIONE-containing cells, *i*.*e*. untransdifferentiated reactive astrocytes or transdifferentiated neurons. The split-Cre system did not show any leakiness in the NCre or CCre only condition (Figure S2). With these results, we designed TRANsCre-DIONE using *Lcn2* and *GFAP* promoters of split-Cre for mouse and *iNOS* and *GFAP* promoters of split-Cre for monkey to express *Neurog2* in reactive astrocytes specifically.

**Figure 1.**
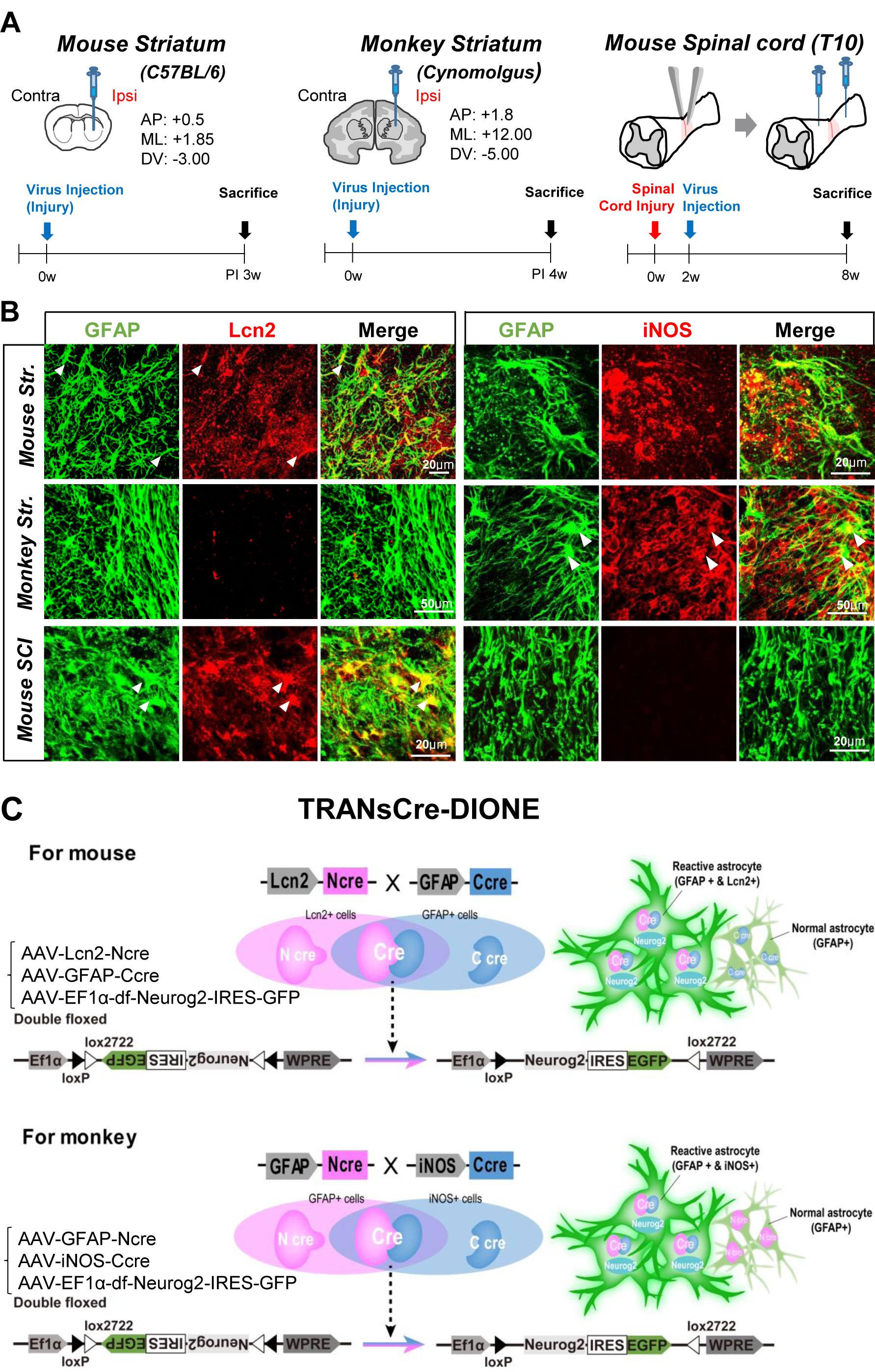
The genetic strategy of transdifferentiation in the brain of a mouse and non-human primate. (A) Experimental protocol for induction of reactiveness by virus injection and the coordination of the surgery in the striatum of the mouse (left), cynomolgus monkey (middle), and the mouse spinal cord (right). Unilateral injection was performed in striatum of mice (AP: +0.5, ML: 1.85, DV: −3.00) and monkey (AP: +1.8, ML: 12.00, DV: −5.00). Injury was given at the level of Thoracic 10 of mouse spinal cord with self-closing forceps. Virus was injected into two point beside the lesion site 2 weeks after the injury. Mice were sacrificed post-injection (PI) 3 weeks. Monkeys were sacrificed PI 4weeks. SCI mice were sacrificed post-injury 8weeks (PI 6 weeks). (B) Lcn2 (red, left panel) and iNOS (red, right panel) expression and co-localization with GFAP (green, both) in the injured striatum (Str.) of mouse and monkey and mouse SCI. (Arrowhead: co-localization of GFAP and Lcn2, or GFAP and iNOS). Scale bar= 20 μm (mouse Str. and mouse SCI), 50 μm (monkey Str.). (C) TRANsCre-DIONE, the strategy for transdifferentiation by expression of *Neurog2* selectively in reactive astrocytes of mouse brain (upper) and monkey brain (lower).

### Transdifferentiation of reactive astrocytes into functional neurons in the mouse brain

To examine whether it is possible that the reactive astrocytes can specifically transdifferentiate into neurons with one transcription factor *in vivo* and to confirm the efficiency of transdifferentiation, we injected TRANsCre-DIONE, a mixture of packaged AAV (1:1:1) for control group (AAV-EF1α-DIO-EGFP with AAV-Lcn2-NCre and AAV-GFAP-CCre) and Neurog2 group (AAV-EF1α-DIO-Neurog2-IRES-EGFP with AAV-Lcn2-NCre and AAV-GFAP-CCre) bilaterally into the mouse striatum (ML 1.85mm, AP 0.5mm, DV −3.0mm). The mice were sacrificed 3 weeks after injection (Figure 2A). We immunostained the tissues with antibodies against GFAP and NeuN for distinguishing the cell types of GFP expressing cells. The efficiency of transdifferentiation was calculated by counting the number of GFAP and NeuN positive cells in GFP expressing cells. Most of the GFP expressing cells (62%, 84 out of 135 cells) were NeuN positive in the Neurog2 group while only 1% (1 out of 93 cells) of GFP expressing cells were NeuN positive in the control group (Figure 2B and 2C). Most of the GFP expressing cells (89%, 83 out of 93 cells) in the control group remained GFAP-positive astrocytes while 28% (39 out of 135 cells) of the GFP expressing cells in Neurog2 group were GFAP positive (Figure 2B and 2C). These results demonstrate that GFAP positive reactive astrocytes transdifferentiate into neurons in the mouse striatum by expressing *Neurog2* solely.

**Figure 2.**
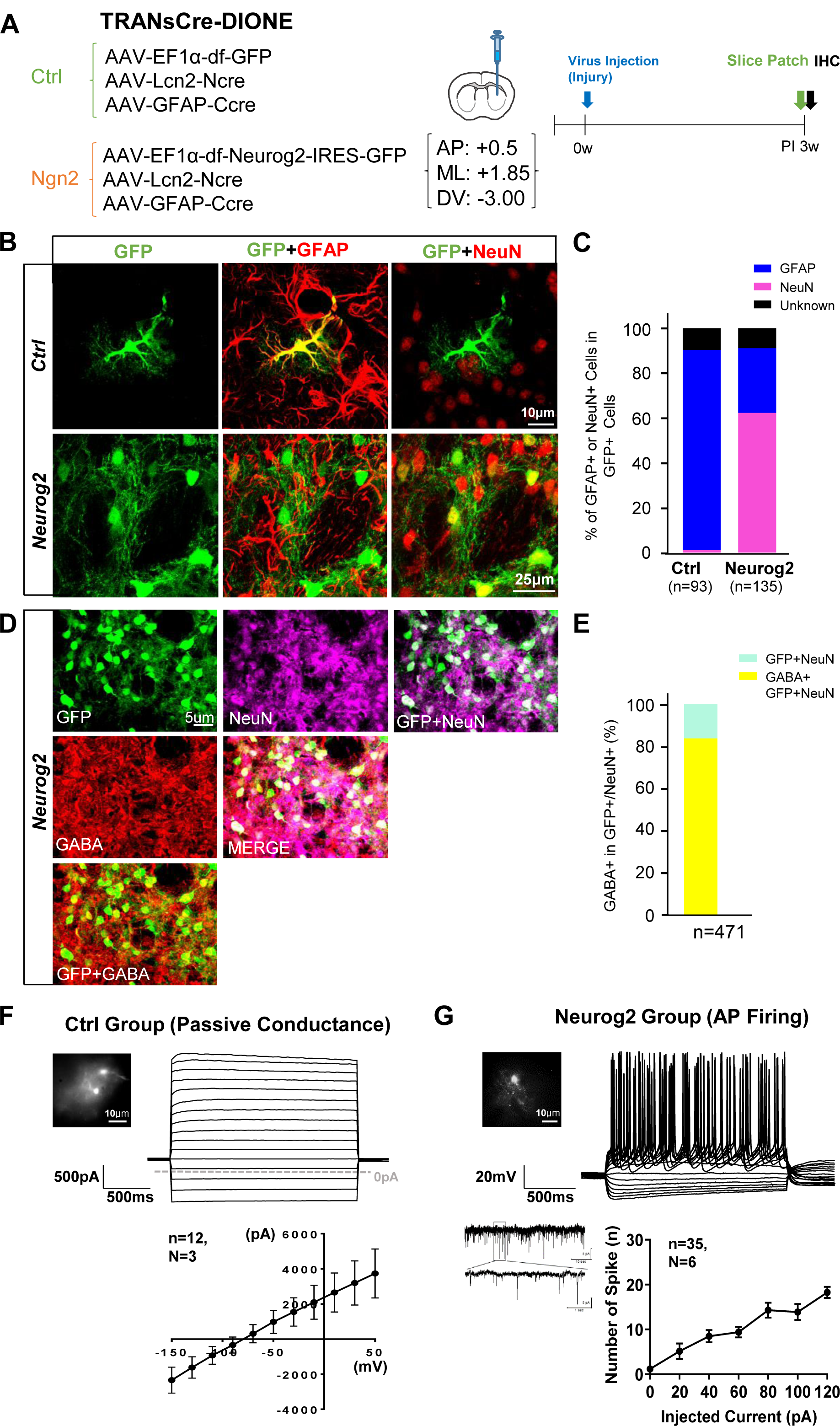
Transdifferentiation of reactive astrocytes into functional neurons in the mouse brain. (A) A mixture of 3 viruses (TRANsCre-DIONE, 2μl of mixture) was injected into the striatum (AP: +0.5, ML: +1.85, DV: −3.0; bilaterally). The mice were sacrificed PI 3 weeks for immunohistochemistry (IHC) and *ex vivo* patch-clamp recording. (B) Colocalization of GFAP (red, middle column) and NeuN (red, right column) with GFP fluorescence (green, left column) in Ctrl (upper) and Neurog2 (lower) group. Scale bar=10 μm (Ctrl group), 25 μm (Neurog2 group). (C) Stacked bar graph showing the percentage of GFAP (blue) and NeuN (pink) positive cells in GFP expressing cells (black: undefined cells). n=93 (Ctrl group), n=135 (Neurog2 group). (D) Colocalization of GABA (red, middle column) and NeuN (magenta, upper) with GFP fluorescence (green) in Neurog2 group. Scale bar= 5 μm. (E) Stacked bar graph showing the number and percentage of GABA positive (yellow) cells in NeuN positive and GFP expressing (cyan) cells. Total n=471 in Neurog2 group. (F) Representative trace of passive conductance (Upper) recorded from GFP expressing cells (inset, scale bar=10 μm) in the control group by voltage-clamp step protocol. Scale bar=500pA, 500ms. Averaged amplitude reached to around 4000pA at 50mV. Averaged I-V graph (lower) from voltage of −150mV to 50mV. N=3, n=12. (G) Representative action potential firing (Upper) and the trace of spontaneous EPSCs (lower, left) from GFP expressing cells (inset, scale bar= 10 μm) in the Neurog2 group. The average of the number of spikes (lower, right) was increased by injected current. N=6, n=35.

Striatal neurons are composed of 100% GABA expressing neurons with 95% medium-spiny neurons expressing both GABA and DARPP32 and 5% of the aspiny GABAergic interneurons (Tepper et al., 2010). In contrast, it has been reported that the transdifferentiated neurons from cultured astrocytes upon over-expression of *Neurog2* were mostly glutamatergic neurons (Heinrich et al., 2010), raising a strong possibility that the transdifferentiated neurons in the striatum are glutamatergic, rather than GABAergic. To determine whether the transdifferentiated neurons in the striatum are GABAergic, we performed double-immunostaining with antibodies against NeuN and GABA (Figure 2D). By surprise, we found that 83.64 % of the NeuN+ GFP-expressing cells were GABA+ (Figure 2E), indicating that the majority of the transdifferentiated neurons were GABAergic in the striatum. These results suggest that over-expression of *Neurog2* causes transdifferentiation into GABAergic neurons in the specific environment surrounded by GABAergic neurons.

To test the functional property of transdifferentiated neurons from reactive astrocytes by forced *Neurog2* expression, we performed whole-cell patch-clamp recording in the infected cells. GFP expressing cells (Figure 2F, inset) in the control group showed a large and linear passive conductance current (Figure 2F, upper panel), which is a unique physiological property of astrocytes in the striatum (Figure 2F). Contrary to the control group, GFP expressing cells (Figure 2F, inset) from the Neurog2 group showed robust action potential firing (Figure 2G, upper panel) by the current-clamp recording with whole-cell patch clamp *ex vivo*. The spontaneous excitatory postsynaptic currents (EPSCs) (Figure 2G, inset) and large sodium and potassium currents were also observed (Figure S4), indicating the maturity of the transdifferentiated neurons (Figure 2G). The similar morphological change and action potential firing were observed within 14 days after transducing *Neurog2* in culture (Figure S3A-C). These results demonstrate that reactive astrocytes transdifferentiate into functional neurons by expressing solely *Neurog2* in reactive astrocytes.

### Transdifferentiation of reactive astrocytes into functional neurons in the monkey’s brain

To investigate if transdifferentiation occurs in the brain of non-human primates, we injected TRANsCre-DIONE into the putamen of cynomolgus monkeys *(Macaca fascularis)* as previously described (An et al., 2016) and harvested the brain tissues after four weeks (Figure 3A). As mentioned above (Figure 1D), *iNOS* was used as a promoter instead of *Lcn2* with *GFAP* for transducing *Neurog2* into reactive astrocytes in the monkey’s brain (Figure 3A). Harvested brain tissues were immunostained with antibodies against GFAP and NeuN as well (Figure 3B). GFP-expressing cells rarely colocalized with NeuN in the control group, while considerable GFP-expressing cells were NeuN positive in the Neurog2 group (Figure 3B). These results demonstrate that the reactive astrocytes in the monkey’s brain also transdifferentiate into NeuN-positive neurons by expressing *Neurog2*. To investigate whether the infected cells completely transdifferentiate into neurons or not, we generated a scatter plot of GFAP intensity *versus* NeuN intensity of all GFP-expressing cells. GFP-expressing cells in the control group (Figure 3C, blue dots) show low NeuN and high GFAP intensity forming a distinct cluster in the left upper quadrant (G+/N-) (Figure 3C). GFP-expressing cells in the Neurog2 group (Figure 3C, pink dots) switched to the right direction in general, indicating that the cells express stronger NeuN signal than in the control group. The cells in the right lower quadrant (G-/N+) might be completely transdifferentiated into neurons, whereas the cells in the right upper quadrant (G+/N+) are in the transition from astrocytes to neurons (Figure 3C). In the Neurog2 group, GFAP-negative (G-) cells occupied almost half of the GFP-expressing cells. Contrary to the Neurog2 group, almost all the GFP-expressing cells in the control group were GFAP-positive (G+) (Figure 3D). While GFP-expressing cells exhibited strong astrocyte properties with high GFAP intensity and low NeuN intensity in the control group (Figure 3E), the cells in the Neurog2 group exhibited much more neuron-like properties with increased NeuN and decreased GFAP intensities, reaching to the similar level of intensity to each other (Figure 3F). In other words, GFP-expressing cells in the Neurog2 group expressed higher NeuN (Figure 3G) and lower GFAP intensity than those of the control group (Figure 3F). Taken together, the reactive astrocytes transdifferentiated into more neuron-like cells in the brain of non-human primates by sole expression of *Neurog2*.

**Figure 3.**
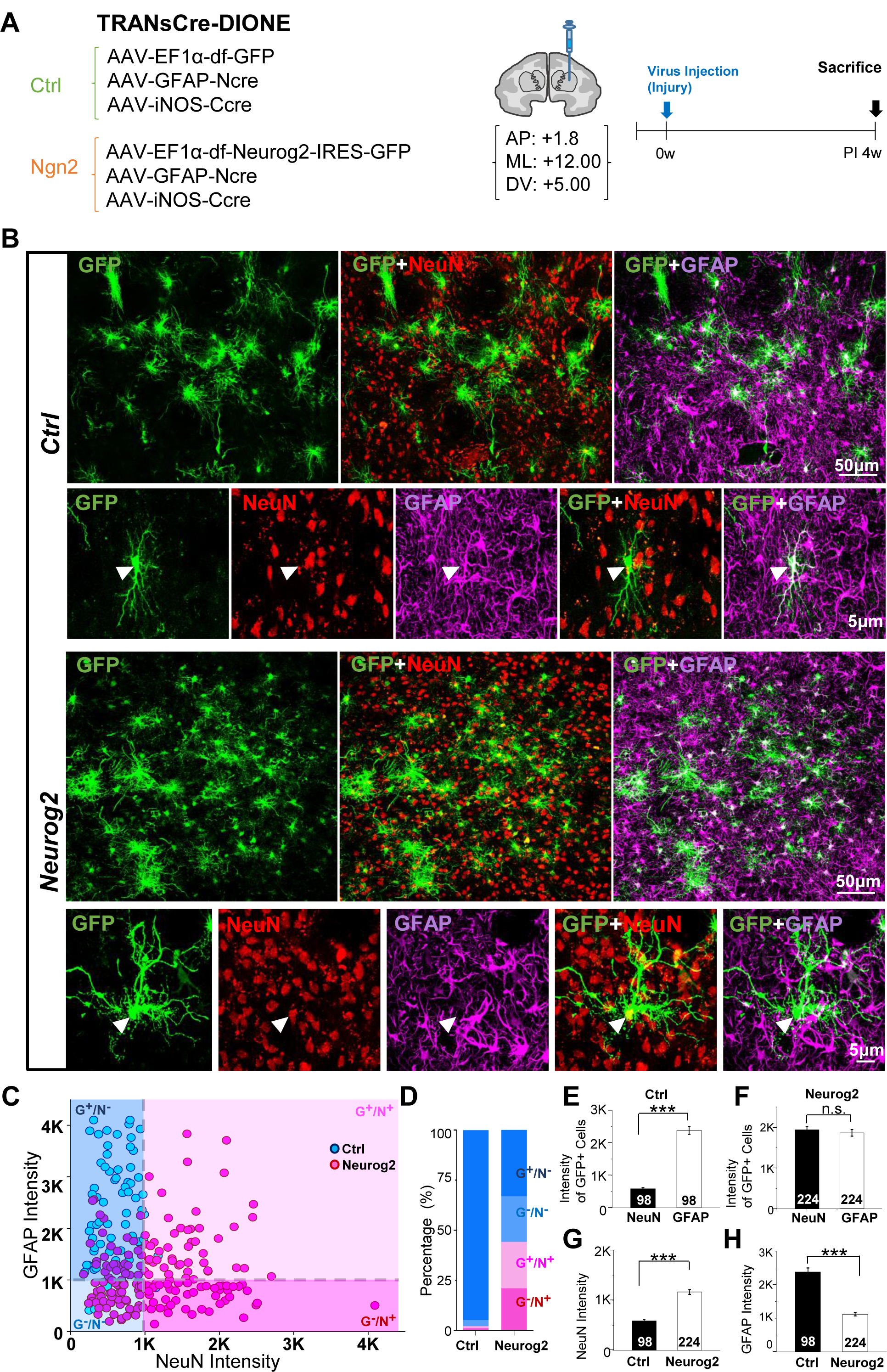
Transdifferentiation of reactive astrocytes into functional neurons in the monkey’s brain. (A) A mixture of 3 viruses (TRANsCre-DIONE, 25μl of the mixture) was injected into the striatum (AP: +1.8, ML: +12.00, DV: −5.0; unilaterally). The monkeys were sacrificed PI 4 weeks for IHC. (B) Colocalization of GFAP (magenta) and NeuN (red) signal in GFP-expressing cells (green) in the Control group (Upper) and Neurog2 group (lower). Scale bar=50 μm (low magnification), 5 μm (magnified images). Arrowhead means the colocalization of GFP and GFAP in Ctrl group, or GFP and NeuN in Neurog2 group. (C) Scatter plot of all GFP expressing cells in Ctrl (blue dot) and Neurog2 (pink dot) group classified as GFAP and NeuN intensity. Quadrant 1(light pink); GFAP+/NeuN+ (G+/N+), Quadrant 2(blue); GFAP+/NeuN- (G+/N-), Quadrant 3(light blue); GFAP-/NeuN- (G-/N-), Quadrant 4(pink); GFAP-/NeuN+(G-/N+) (Reference lines are drawn for dividing + or – at 1K of both intensities.). n=98 (Ctrl group), n=224 (Neurog2 group). (D) Stacked bar graph of the percentage of G+/N-, G-/N-, G+/N+ and G-/N+ in Ctrl and Neurog2 groups. (E) Comparison of NeuN and GFAP intensity of GFP-expressing cells in Ctrl group. Unpaired student t-test (n.s.=p>0.05, *p<0.05, **p<0.01, ***p<0.001, ****p<0.0001). n=98 (Ctrl group). (F) Comparison of NeuN and GFAP intensity of GFP+ cells in Neurog2 group. Unpaired student t-test. n=224 (Neurog2 group). (G) Comparison of NeuN intensity of GFP+ cells in Ctrl and Neurog2 group. Unpaired student t-test. n=98 (Ctrl group), n=224 (Neurog2 group). (H) Comparison of GFAP intensity of GFP+ cells in the Ctrl and Neurog2 group. Unpaired student t-test. n=98 (Ctrl group), n=224 (Neurog2 group).

### Transdifferentiation of scar-forming reactive astrocytes into functional neurons in the spinal cord injury mouse model

To investigate whether the transdifferentiated neurons from reactive astrocytes can ameliorate the motor impairment caused by spinal cord injury, we generated the spinal cord injury mouse model with forceps compressing of spinal cord at Thoracic vertebra (T10) (Figure 4B) and injected TRANsCre-DIONE (AAV-EF1α-DIO-Neurog2-IRES-EGFP with AAV-Lcn2-NCre and AAV-GFAP-CCre) into 1mm above and below the injury site two weeks after injury (Figure 4A and 4B). Starting from the time of injury, every week we measured the Basso Mouse Scale (BMS) that represents locomotion defects during recovery. SCI mice were scored from zero to nine based on each mouse’s ability to move its lower body. The impaired movement was significantly ameliorated to an average BMS score of 3.88 at 8 week in the SCI/Neurog2 group, while the mice in SCI/Ctrl and SCI/PBS group were scored below an average of 1 (Figure 4C). These results indicate that TRANsCre-DIONE can alleviate the motor impairment of SCI mouse model. To investigate whether the transdifferentiated cells display functional properties of neurons, GFP-expressing cell (inset, Figure 4D) in the Neurog2 group was recorded by whole-cell patch clamp. GFP-expressing cell showed robust action potential firing (Figure 4D), large sodium currents (Figure 4E), and numerous spontaneous EPSCs (Figure 4F). Taken together, TRANsCre-DIONE causes an improvement of motor behavior and transdifferentiation into functional neurons in SCI.

**Figure 4.**
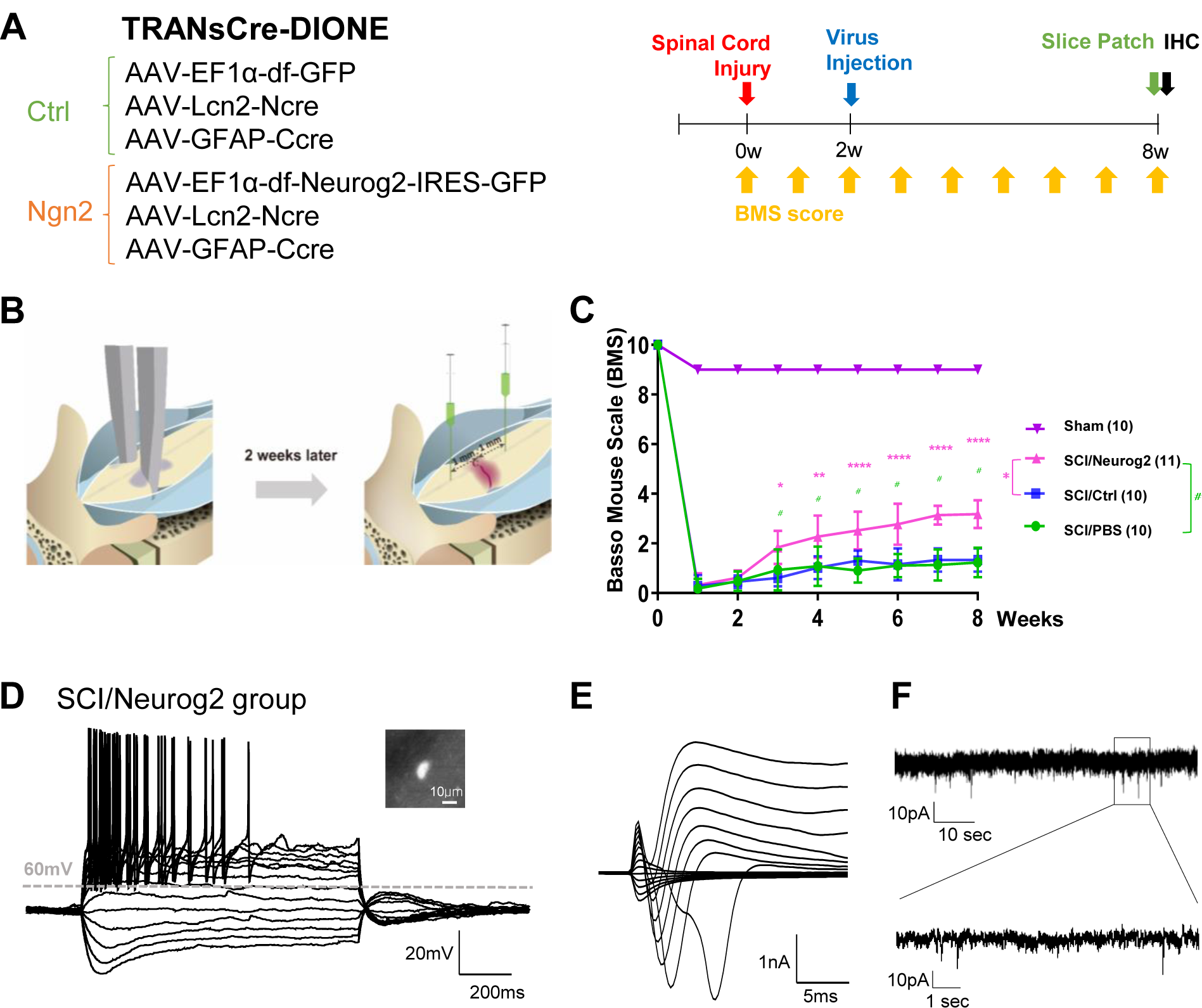
Transdifferentiation of scar-forming reactive astrocytes into functional neurons in the spinal cord injury mouse model. (A) A mixture of 3 viruses (TRANsCre-DIONE, 1μl of mixture) was injected into the spinal cord. Basso Mouse Scale (BMS) score was measured in every week from the first week of the injury to the last week. SCI mice were sacrificed at 8 weeks after injury (PI 6 weeks) for IHC and *ex vivo* patch-clamp recording. (B Schematic diagram of the protocol for setting the SCI model and virus injection in the spinal cord. The SCI mouse model was set by forceps compressing protocol (compression for 3 sec, at Thoracic10). A mixture of 3 viruses (TRANsCre-DIONE, 1μl of mixture, upper and lower of 1mm of the injury site) was injected into the spinal cord 2 weeks after injury. (C) BMS scores of sham, SCI/Neurog2, SCI/Ctrl and SCI/PBS groups. BMS score was measured in every week for following the functional recovery of the SCI model. (magenta; Sham group, pink; SCI/Neurog2 group, blue; SCI/Ctrl group, green; SCI/PBS group). Dunnett’s multiple comparison test, with individual variances computed for each comparison (#p<0.05). n=10 (sham group), 11 (SCI/Neurog2 group), 10 (SCI/Ctrl group), 10 (SCI/PBS group). (D Action potential firing was in GFP-expressing cell (inset, scale bar=10 μm) in Neurog2 group. Scale bar= 20mV, 200ms. (E) Large sodium channel current was shown in voltage-clamp recording from GFP-expressing cell in the Neurog2 group. Scale bar= 1nA, 5ms. (F) Spontaneous EPSCs from GFP-expressing cell in the Neurog2 group. Scale bar= 10pA, 10sec (upper); 10pA, 1sec (enlarged trace).

### Tissue recovery in SCI via transdifferentiation of scar-forming reactive astrocytes into neurons

To test whether over-expression of *Neurog2* in reactive astrocytes causes tissue recovery, we performed Erich-Chrome (EC) staining with the spinal cord tissues to measure the myelinated, gray matter and total areas (Figure 5A). Although the tissues were severely shrunken and damaged in SCI/PBS group and SCI/Ctrl group, tissues in Neurog2 group were recovered almost to the sham group (Figure 5A). The gray matter, myelinated and total areas were decreased in SCI/PBS (27.95%, 28.77%, 28.39%) and SCI/Ctrl (37.74%, 12.69%, 26.01%) groups compared to the sham group, and the areas were significantly increased in SCI/Neurog2 group (66.37%, 49.51%, 57.45%) (Figure 5B). To explore whether the damaged axon and dendrites were recovered in the injured spinal cord, we immunostained the longitudinal spinal cord sections with an antibody against MAP2. Although the MAP2 signal was almost disappeared around the injury site in SCI/PBS and SCI/Ctrl groups, it was remarkably recovered in SCI/Neurog2 group (Figure 5C). These results demonstrate that myelinated axon and the damaged tissue structure might be restored via TRANsCre-DIONE.

**Figure 5.**
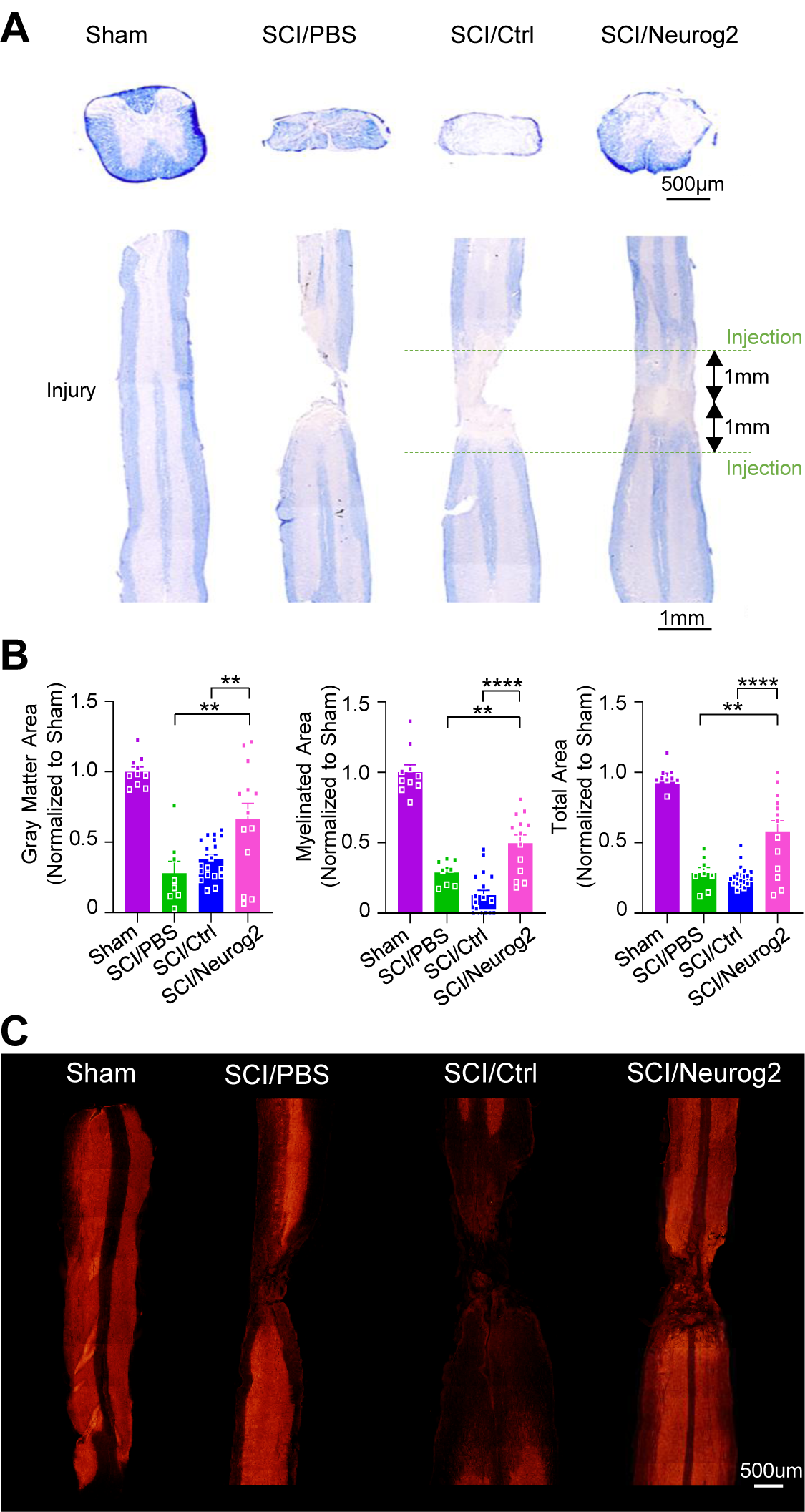
Tissue recovery in SCI via transdifferentiation of scar-forming reactive astrocytes into neurons. (A) Eriochrome Cyanine (EC) staining for myelin staining in cross and longitudinal sections in sham, SCI/PBS, SCI/Ctrl and SCI/Neurog2 groups. (Black dot line; injury site, green dot line; injection site). Scale bar= 500 μm (cross section), 1mm (longitudinal section). (B) Scattered bar graph showing the area of gray matter (left), myelinated area (middle), and total area (right) in sham (magenta), SCI/PBS (green), SCI/Ctrl (blue), SCI/Neurog2 (pink) groups (Each value was normalized by sham group). Unpaired non-parametric one-way ANOVA with Kruskal-Wallis test. n= 10 (sham group), 8 (SCI/ PBS group), 19 (SCI/Ctrl group), 13 (SCI/Neurog2 group). (C) MAP2 staining of longitudinal sections of Sham, SCI/PBS, SCI/Ctrl, and SCI/Neurog2 groups. (Scale bar= 500μm, 8 weeks)

### Transdifferentiation of reactive astrocytes into motor neurons in SCI

Isl1 is necessary for differentiating most of the subtypes of motor neurons such as VChAT+, ChAT+, CHT+ neurons, and Isl1-expressing cholinergic motor neurons are major cell types in the spinal cord (Cho et al., 2014). To examine whether over-expression of *Neurog2* induces transdifferentiation of reactive astrocytes into specific neurons, we immunostained the spinal cord tissues with antibodies against Isl1, the key molecular marker for motor neuron (Tang et al., 2017), and other neuronal markers such as MAP2 and NeuN. MAP2 signal was disappeared in SCI/PBS and SCI/Ctrl groups and significantly restored in SCI/Neurog2 group (Figure 6A and 6C). MAP2 signal in GFP-expressing cells was also significantly increased in SCI/Neurog2 group compared to SCI/Ctrl group (Figure 6A and Figure6D). 97.32% of the GFP-expressing cells were MAP2 positive in SCI/Neurog2 group, while 98.88% of the GFP-expressing cells were GFAP positive in SCI/Ctrl group. These results indicate that the most of the GFP expressing cells were transdifferentiated into MAP2+ neurons via expression of *Neurog2* in SCI.

NeuN signal was also disappeared in SCI/PBS and SCI/Ctrl groups and significantly recovered in SCI/Neurog2 group (Figure 6B and Figure 6H). NeuN signal in GFP-expressing cells was also significantly increased in SCI/Neurog2 group compared to SCI/Ctrl group (Figure 6B and Figure 6I). To determine the transdifferentiation efficiency and specificity, we performed detailed analysis of GFP-expressing cells by plotting NeuN versus GFAP (Figure S4B and S4C). We found that 87% (67 out of 77 cells) of GFP-expressing cells were NeuN positive in SCI/Neurog2 group, while only 3.92% (4 out of 102 cells) were NeuN positive in SCI/Ctrl group (Figure 6F). Finally, the Isl1 signal was disappeared in SCI/PBS and SCI/Ctrl groups and significantly increased in SCI/Neurog2 group (Figure 6B and 6J). The Isl1 signal in GFP-expressing cells was also significantly increased in SCI/Neurog2 group compared to SCI/Ctrl group (Figure 6B and Figure 6K). The percentage of Isl1+/GFP+ motor neurons in total Isl1+ motor neurons was significantly increased in SCI/Neurog2 group, while all the Isl1+ motor neurons were GFP-in SCI/Ctrl group (Figure 6B and Figure 6L). These results indicate that reactive astrocytes transdifferentiated into Isl1+ motor neurons in the injured spinal cord by over-expression of *Neurog2*. To explore whether over-expression of *Neurog2* in reactive astrocytes affects the reactivity of astrocytes, we immunostained the tissues with the antibody against GFAP. While the GFAP signal was increased in SCI/PBS and SCI/Ctrl groups, GFAP was significantly decreased in SCI/Neurog2 group (Figure 6A, 6B and 6G). Taken together, TRANsCre-DIONE induces transdifferentiation of reactive astrocytes into motor neurons and attenuates the reactivity of astrocytes in the injured spinal cord.

**Figure 6.**
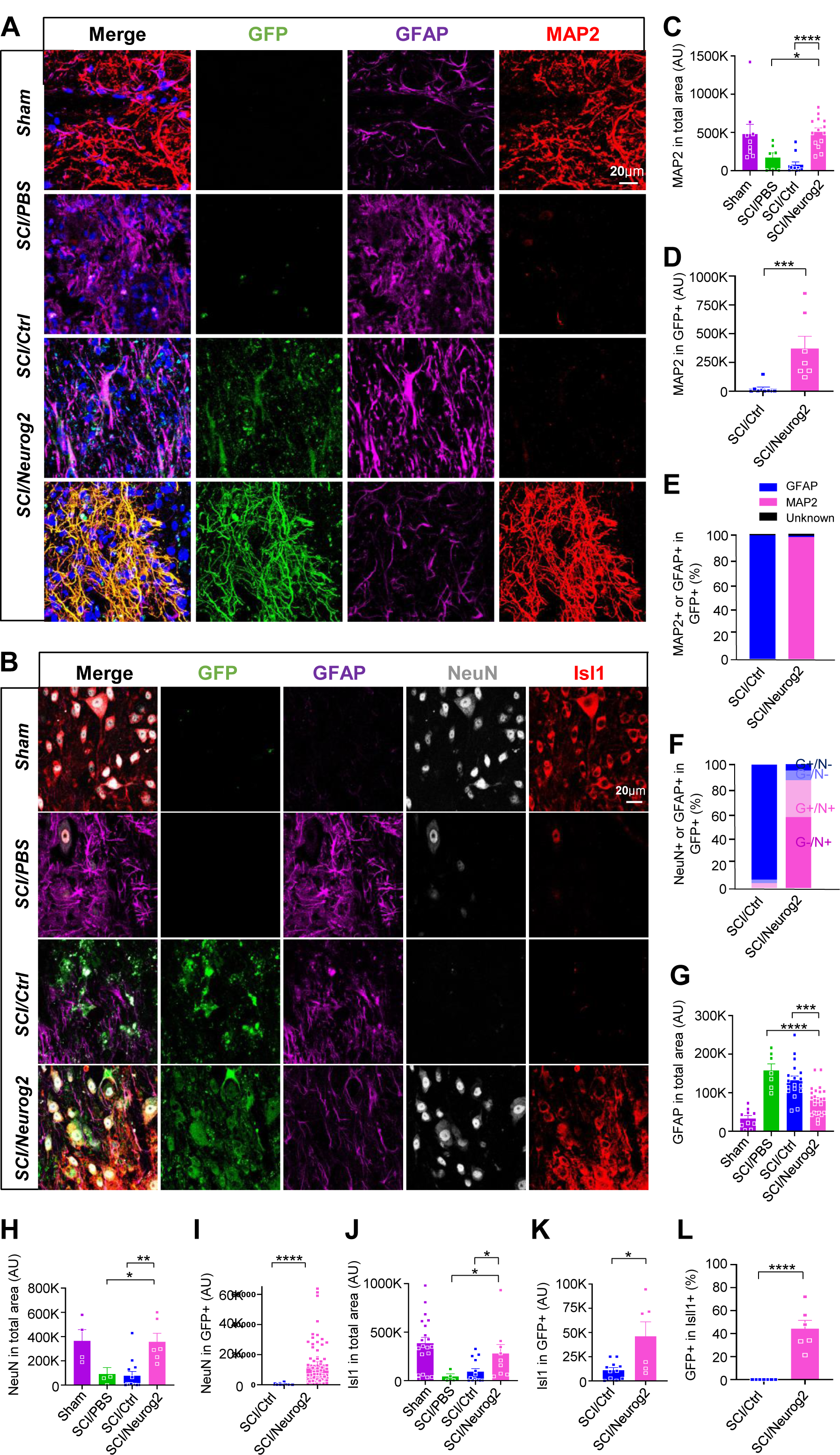
Transdifferentiation of reactive astrocytes into motor neurons in SCI. (A) Colocalization of GFAP (magenta) and MAP2 (red) with GFP fluorescence (green) in sham, SCI/PBS, SCI/Ctrl and SCI/Neurog2 groups. Scale bar=20 μm. (B) Colocalization of GFAP (magenta), NeuN (gray) and Isl1 (red) with GFP fluorescence (green) in sham, SCI/PBS, SCI/Ctrl and SCI/Neurog2 groups. Scale bar=20 μm. (C) Scattered bar graph showing comparison of MAP2 intensity in the total area. Multiple comparison for unpaired non-parametric one-way ANOVA with Kruskal-Wallis test. n=9 (sham group), 7 (SCI/PBS group), 11 (SCI/Ctrl group), 14 (SCI/Neurog2 group). (D) Scattered bar graph showing comparison of MAP2 intensity in GFP-expressing cells in SCI/Ctrl and SCI/Neurog2 groups. Unpaired non-parametric t-test with Mann-Whitney test. n=11 (SCI/Ctrl group), 14 (SCI/Neurog2 group). (E) Stacked bar graph showing the proportion of GFAP (blue) or MAP2 (pink) positive cells in GFP expressing cells. (Undefined: black). (F) Stacked bar graph showing the proportion of GFAP and NeuN positive or negative cells in GFP expressing cells. G+/N- (blue, quadrant 2 of Figure S4C), G-/N- (light blue, quadrant 3 of Figure S4C), G+/N+ (light pink, quadrant 1 of Figure S4C) and G-/N+ (pink, quadrant 4 of Figure S4C). n=102 in SCI/Ctrl, n=77 in SCI/Neurog2 group. (G) Scattered bar graph showing comparison of GFAP intensity in the total area. Multiple comparison for unpaired non-parametric one-way ANOVA with Kruskal-Wallis test. n=10 (sham group), 8 (SCI/PBS group), 18 (SCI/Ctrl group), 24 (SCI/Neurog2 group). (H) Scattered bar graph showing comparison of NeuN intensity in the total area. Multiple comparison for unpaired non-parametric one-way ANOVA with Kruskal-Wallis test. n=4 (sham group), 4 (SCI/PBS group), 13 (SCI/Ctrl group), 6 (SCI/Neurog2 group). (I) Scattered bar graph showing comparison of NeuN intensity in GFP-expressing cells in SCI/Ctrl and SCI/Neurog2 groups. Unpaired non-parametric t-test with Mann-Whitney test. n=66 (SCI/Ctrl group), 65 (SCI/Neurog2 group). (J) Scattered bar graph showing comparison of Isl1 intensity in the total area. Multiple comparison for unpaired non-parametric one-way ANOVA with Kruskal-Wallis test. n=20 (sham group), 4 (SCI/PBS group), 14 (SCI/Ctrl group), 9 (SCI/Neurog2 group). (K) Scattered bar graph showing comparison of Isl1 intensity in GFP-expressing cells in SCI/Ctrl and SCI/Neurog2 groups. Unpaired non-parametric t-test with Mann-Whitney test. n=11 (SCI/Ctrl group), 6 (SCI/Neurog2 group). (L) Scattered bar graph showing comparison of GFP-expressing cells in Isl1-positive cells in SCI/Ctrl and SCI/Neurog2 groups. Unpaired non-parametric t-test with Mann-Whitney test. n=7 (SCI/Ctrl group), 6 (SCI/Neurog2 group).

### Transdifferentiated neurons are mostly originated from non-proliferating cells

To investigate whether the transdifferentiated neurons are derived from proliferating cells, we performed labeling with Bromodeoxyuridine (BrdU) which is incorporated into DNA during cell proliferation (Russo et al., 1984). BrdU was administered once a day for one week to the SCI mice before they were sacrificed at the time of 3-week (Figure S5B) and 8-week (Figure7B). For detecting BrdU signals in the specific cell types from GFP-expressing cells, we double-stained the tissues with antibodies against GFAP or NeuN or MBP (myelination binding protein), the cell markers of astrocyte, neuron and oligodendrocyte, respectively (Figure 7A and Figure S5A). In SCI/Ctrl group, only 12.06% of the GFP+ cells and 9.60% of the GFP+/GFAP+ cells were labeled with BrdU, while most of the GFP-cells were labeled with BrdU (87.90%, Figure 7C). Similarly, in SCI/Neurog2 group, only 5.90% of the GFP+ cells and 1.18% of the GFP+/GFAP+ cells were labeled with BrdU, while almost all of the GFP-cells were labeled with BrdU (94.09%, Figure 7C). These results indicate that most of the GFP+ cells are not proliferating in the late stage (Figure 7C), as well as in the early stage of transdifferentiation (Figure S5C). Double-labeling with BrdU and NeuN resulted in BrdU labeling of only 10.36% of the GFP+ cells and 0.39% of the GFP+/NeuN+ cells in SCI/Ctrl group. Similarly, BrdU labeled only 9.82% of the GFP+ cells and 9.62% of the GFP+/NeuN+ cells in SCI/Neurog2 group (Figure 7D). In contrast, most of the GFP-cells in SCI/Ctrl group (89%) and in SCI/Neurog2 group (85%) were labeled with BrdU (Figure 7D). These results indicate that most of the transdifferentiated GFP+/NeuN+ cells are not proliferating in the late stage (Figure 7A and 7D), as well as in the early stage of the transdifferentiation (Figure S5D). There were no GFP+/MBP+ cells in both SCI/Ctrl and SCI/Neurog2 groups, indicating that oligodendrocytes were not transdifferentiating into neurons (Figure 7E). Taken together, these results demonstrate that GFP+ transdifferentiated neurons are mostly originated from BrdU negative non-proliferating cells in the early stage as well as in the late stage of transdifferentiation.

**Figure 7.**
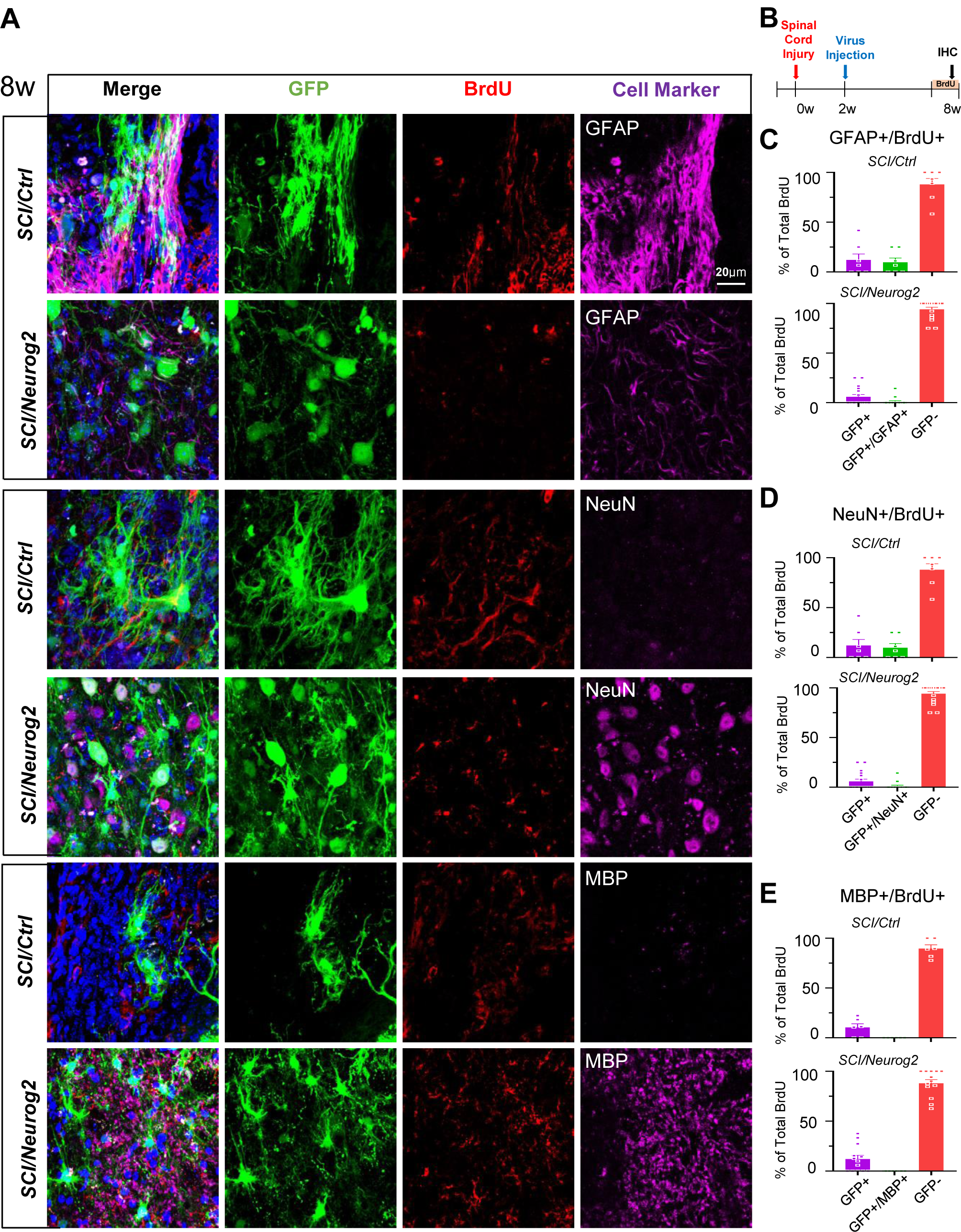
Transdifferentiated neurons are mostly originated from non-proliferating cells. (A) Colocalization of BrdU (red, treated for 1week from 7 weeks after injury) in GFP-expressing cells (green) with cell markers (magenta; GFAP, upper; NeuN, middle; MBP, lower) in SCI/Ctrl and SCI/Neurog2 groups. Scale bar=20 μm. (B) Timeline of treating BrdU and IHC after sacrificing in SCI mice. (C) Scattered bar graph showing comparison of percentage of BrdU-positive cells in GFP+, GFAP+/GFP+ and GFP-cells in SCI/Ctrl and SCI/Neurog2 groups. n=7 (SCI/Ctrl group), 17 (SCI/Neurog2 group). (D) Scattered bar graph showing comparison of percentage of BrdU-positive cells in GFP+, NeuN+/GFP+ and GFP-cells in SCI/Ctrl and SCI/Neurog2 groups. n=7 (SCI/Ctrl group), 17 (SCI/Neurog2 group). (E) Scattered bar graph showing comparison of percentage of BrdU-positive cells in GFP+, MBP+/GFP+ and GFP-cells in SCI/Ctrl and SCI/Neurog2 groups. n=7 (SCI/Ctrl group), 17 (SCI/Neurog2 group).

To investigate whether the reduction of reactive astrocytes influences the proliferation of surrounding cells (BrdU+/GFP-) near the transdifferentiated neurons, we assessed the proportion of BrdU+ cells in each cell type (GFAP+, NeuN+, MBP+) among the surrounding GFP-cells (Table 1). We found that the proportion of proliferating astrocytes near the transdifferentiated cells (GFP-/BrdU+/GFAP+) were decreased from 9.62% (SCI/Ctrl) to 4.55% (SCI/Neurog2) in the early stage (3 week, Figure S5A), from 12.50% (SCI/Ctrl) to 10.20% (SCI/Neurog2) in the late stage (8 week, Figure 7A). In contrast, the proportion of GFP-/BrdU+/NeuN+ cells was increased from 0.00% (SCI/Ctrl) to 4.49% (SCI/Neurog2) in the early stage (3 week, Figure S5A), from 1.93% to 9.80% in the late stage of transdifferentiation (8 week, Figure 7A). Consistently, the proportion of GFP-/BrdU+/MBP+ cells was increased from 9.13% (SCI/Ctrl) to 12.03% (SCI/Neurog2) in the early stage (3 week, Figure S5A), from 1.33% to 11.94% in the late stage of transdifferentiation (8 week, Figure 7A). Taken together, these results implicate that the regeneration of neighboring neurons (GFP-/BrdU+/NeuN+) and oligodendrocytes (GFP-/BrdU+/MBP+) was enhanced when the astrocytic reactivity was decreased by TRANsCre-DIONE in SCI/Neurog2 group in both early and late stages.

**Table 1.** Proportion of non-transdifferentiated (GFP-) proliferative (BrdU+) astrocytes (GFAP+), neurons (NeuN+) and oligodendrocytes (MBP+)

|  |  | GFAP |  | NeuN |  | MBP |  |
| --- | --- | --- | --- | --- | --- | --- | --- |
|  |  | SCI/Ctrl | SCI/Neurog2 | SCI/Ctrl | SCI/Neurog2 | SCI/Ctrl | SCI/Neurog2 |
| 3 week | Total BrdU+ | 52 | 22 | 138 | 156 | 230 | 158 |
|  | GFP-/BrdU+/Marker+ | 5 | 1 | 0 | 7 | 21 | 19 |
|  |  | 9.62% | 4.55% | 0.00% | 4.49% | 9.13% | 12.03% |
| 8 week | Total BrdU+ | 64 | 98 | 207 | 204 | 75 | 134 |
|  | GFP-/BrdU+/Marker+ | 8 | 10 | 4 | 20 | 1 | 16 |
|  |  | 12.50% | 10.20% | 1.93% | 9.80% | 1.33% | 11.94% |

## DISCUSSION

In this study, we have developed and characterized a novel molecular tool set, TRANsCre-DIONE, which is highly effective (62% in mouse striatum, 87% in SCI) in replacing dead neurons in severe injury conditions (Figure S6). The highlights of our study are that 1) TRANsCre-DIONE converts reactive astrocytes into neurons by a sole over-expression of *Neurog2*, 2) reactive astrocytes are specifically targeted using split-Cre under two promoters, *GFAP* and *Lcn2* in mouse and *GFAP* and *iNOS* in monkey, 3) TRANsCre-DIONE reduces astrogliosis, replaces dead neurons and alleviates symptoms of SCI, and 4) the transdifferentiated-neurons are GABA+ medium spiny neurons in the striatum and Isl1+ motor neurons in the spinal cord. TRANsCre-DIONE shows an excellent therapeutic potential in human as the combination of *GFAP* and *iNOS* was highly effective in the striatum of non-human primate. TRANsCre-DIONE not only induced direct reprograming of the infected cells but also caused an indirect enhancement of regeneration in neighboring cells, possibly due to removing toxic reactive astrocytes.

In our recent studies, we have newly defined ‘reactive’ astrocytes as the astrocytes which respond to damaging stimulations such as traumatic brain injury or toxic protein aggregates to turn on the MAO-B-dependent GABA synthetic pathway as a consequence of degradation of toxic materials (Chun et al., 2018; Chun and Lee, 2018). These ‘reactive’ astrocytes are known to be detrimental to neighboring neurons and can be easily distinguished from ‘active’ astrocytes, which are known to be beneficial, are present in conditions of enriched environment, and show an enhanced expression of proBDNF with no GABA (Chun et al., 2018; Chun and Lee, 2018). Although the active and reactive astrocytes can be easily distinguished by the two markers, proBDNF and GABA, it is not very easy to distinguish them by the morphology, because both the active and reactive astrocytes show hypertrophy, which is simply an increased level of GFAP expression. In other words, it is very difficult to distinguish between beneficial active astrocytes and detrimental reactive astrocytes using GFAP alone. By utilizing the reactive-astrocyte-specific inducible genes such as *Lcn2* and *iNOS* for promoters of split-Cre in combination with *GFAP* promoter, we have overcome the limitations of solely using *GFAP* promoter and succeeded in specifically targeting reactive astrocytes while sparing active and normal astrocytes. Because Lcn2 and iNOS were never observed in the uninjured tissues and they were induced only upon injury (Figure S1), we interpret the presence of the reporter GFP of TRANsCre-DIONE as a direct representation of reactive astrocytes. We have observed that TRANsCre-DIONE shows extremely high specificity for reactive astrocytes, targeting over 90.21% of reactive astrocytes in the injured mouse striatum (Figure 2C), 93.13% in the injured monkey striatum (Figure 3C), and 98.48% in SCI (Figure 6F). It is worth highlighting the near complete specificity of the combination of *iNOS* and *GFAP* promoters in the monkey striatum. Such complete specificity is most likely due to the specificity of iNOS for reactive astrocytes in non-human primate. These results raise a promising therapeutic potential for an immediate clinical application of TRANsCre-DIONE in injured human brain.

The specificity towards reactive astrocytes or neurons can be estimated by calculating the proportion of GFAP+/GFP+ or NeuN+/GFP+ cells in total GFP+ cells in Ctrl group. In other words, the specificity towards NeuN+ neurons in Ctrl group represents the leakiness of TRANsCre-DIONE towards resident neurons. In the Ctrl group, we observed that only 1 out of 93 (1.07%) GFP+ cells was NeuN+ in the mouse striatum (Figure 2C), 1 out of 98 (1.03%) GFP+ cells was NeuN+ in the monkey’s striatum (Figure 3C), and 4 out of 102 (3.92%) GFP+ cells was NeuN+ in the mouse spinal cord (Figure 6F). These results indicate that TRANsCre-DIONE shows virtually no leakiness towards resident neurons. In contrast, previously reported molecular tools display a significant degree of leakiness towards resident neurons. Having a significant leakiness is problematic because it leads to an inadvertent over-estimation of the efficiency of transdifferentiation. For example, in a recent study using GFAP::Cre for selective expression of two transcription factors, *NeuroD1* and *Dlx2*, authors reported that 10% of infected cells were NeuN+ in control condition (Wu et al., 2020): the efficiency of transdifferentiation was 80% in the striatum of R6/2 Huntington disease mouse model. If at least 10% of the targeted cells were already neurons, the reported transdifferentiation efficiency must have been over-estimated. In another study using only GFAP promoter to target reactive astrocytes with retroviral GFAP::NeuroD1-IRES-GFP, the transdifferentiation efficiency was reported to be over 80% (Guo et al., 2014): this efficiency value might have been over-estimated due to a leakiness. Unfortunately, the authors did not report a corresponding value for leakiness. Because TRANsCre-DIONE shows virtually no leakiness towards neurons, our estimation of transdifferentiation efficiency at 62% in the mouse striatum and 87% in SCI (Figure 6F) is considered to be fairly accurate with less likelihood of over-estimation. Therefore, the performance of TRANsCre-DIONE in specificity as well as in efficiency of transdifferentiation is unprecedentedly high.

The high transdifferentiation efficiency of TRANsCre-DIONE can be attributed to the use of DIO system (Tronche et al., 2002) in combination with the strong *EF1α* promoter (Wang et al., 2017). Unlike other previously reported molecular tools utilizing a simple *GFAP* promoter to drive an expression of transcription factors (Guo et al., 2014; Su et al., 2014; Wu et al., 2020), this unique combination ensures a sustained expression of *Neurog2* in the targeted reactive astrocytes even in the transitional and transdifferentiated state. The net effect is that the transdifferentiation efficiency is sustained high throughout the lifetime of transdifferentiated neurons. Furthermore, our approach of *EF1*α-driven *DIO-Neurog2* system safely prevents an inadvertent expression of *Neurog2* in untargeted cells even under the strong universal promoter, *EF1*α. This safety feature ensures no inadvertent expression of *Neurog2* in untargeted normal astrocytes or other cell types. The net effect of this safety feature is manifested as the virtually zero leakiness and high specificity of TRANsCre-DIONE.

Our study reveals that the sole use of the transcription factor, *Neurog2* is an excellent choice for transdifferentiation of reactive astrocytes into neurons. We found that *Neurog2* alone is capable of transdifferentiating reactive astrocytes – in cortical astrocyte culture, mouse and monkey striatum, and mouse spinal cord – into neurons without a need for additional transcription factors. This is in great contrast to the previous studies in which multiple transcription factors – in combination of *NeuroD1* with *DLX2* (Wu, 2020), *Neurog2* with *SOX11* and *Lhx3* (Liu et al., 2016) and *BRN2* with *MYT1L*, and *FEZF2* (Miskinyte et al., 2017) – have been utilized. Use of only one transcription factor provides an important advantage in that, one can minimize the number of viral vectors and genes of interest to deliver to the site of injury. More importantly, *Neurog2* possesses a chameleon-like property that allows transdifferentiation of reactive astrocytes into various neuronal types depending on the surrounding environment and region of the injury site. It has been shown that *Neurog2* can transdifferentiate cultured astrocytes into glutamatergic neurons (Heinrich et al., 2010). It has been also shown that *Neurog2* can transdifferentiate astrocytes into glutamatergic neurons or GABAergic neurons *in vivo* in cortex, cerebellum, and spinal cord, although the majority was glutamatergic (Hu et al., 2019). However, the transdifferentiation efficiency was less than 10% probably due to the use of *GFAP* promoter alone to drive *Neurog2* expression (Hu et al., 2019). We have shown that *Neurog2* under TRANsCre-DIONE can transdifferentiate reactive astrocytes into GABA+ medium spiny neurons at high rate (83.64 %) in the mouse striatum (Figure 2D-E). In SCI, *Neurog2* was capable of transdifferentiating reactive astrocytes into isl1+ cholinergic motor neurons (Figure 6K). These results imply that the same transcription factor *Neurog2* can cause a transdifferentiation of the target cells into different type of neurons and that the environmental factors strongly influence the fate of the *Neurog2*–expressing transdifferentiating cells. How is this possible for *Neurog2*? As an upstream master switch gene, *Neurog2* is known to induce sequential expression of other neurogenesis-related downstream transcription factors such as *NeuroD1, NeuroD4, Hes6*, and *MyT1* (Seo et al., 2007). Thus, *Neurog2* has a potential to induce transdifferentiation into diverse types of neurons by turning on a specific switch depending on the niche of the residential brain regions. Unlike *Neurog2, NeuroD1* appears to transdifferentiate astrocytes into mostly glutamatergic neurons in hippocampus (Roybon et al., 2015; Wu et al., 2020) when it is used solely. *NeuroD1* appears to require other transcription factors such as *DLX2* in combination to transdifferentiate into other type of neurons other than glutamatergic neurons (Liu et al., 2020; Wu, 2020). In support of this idea, it has been recently demonstrated that a gene-silencing of *PTBP1* induces transdifferentiation of Müller glia to retinal ganglion cells in the retina, whereas it induces transdifferentiation of astrocytes into dopaminergic neurons in the striatum without additional factors (Zhou et al., 2020). Interestingly, authors found that there was no significant change in the level of *NeuroD1* in the transdifferentiated retinal ganglion cells or dopaminergic neurons in the striatum, indicating that *NeuroD1* may not be the downstream transcription factor of *PTBP1*-gene-silencing. It will be interesting to test whether *Neurog2* is the downstream transcription factor of *PTBP1*-gene-silencing. Taken together, *Neurog2* shows several advantages over *NeuroD1* and can be considered as the most optimal transcription factor for transdifferentiation of reactive astrocytes into various types of neurons.

In this study, we have achieved multiple goals using TRANsCre-DIONE: elimination of detrimental reactive astrocytes, replacement of dead neurons, and alleviation of motor symptoms in SCI model. It has been proposed that the “scar-forming reactive astrocytes” (defined as “severe reactive astrocytes” in our study) in SCI inhibit axon regeneration under the chronic pathological condition, while “reactive astrocytes” (defined as “mild reactive astrocytes” in our study) serve protective roles in acute pathological condition (Kijima et al., 2019). According to this proposed model, the “reactive astrocytes” can be transformed into normal astrocytes or scar-forming reactive astrocytes by the surrounding environment in the spinal cord (Okada et al., 2018). Thus, we focused on the effect of eliminating the detrimental scar-forming reactive astrocytes to provide a niche for axon regeneration. Consistence with the proposed model, we observed both a significant decrease in GFAP signal in SCI/Neurog2 group, indicating a relief from astrocytic reactivity and a significant increase in MAP2 signal, indicating an enhanced axon regeneration. In addition, TRANsCre-DIONE synergistically increased appearance of the BrdU+ proliferating cells that were GFP- (Table 1), indicating an enhanced regeneration in the surrounding cells. Such unexpected synergistic effects by removing the detrimental reactive astrocytes imply a pro-regenerative effect of TRANsCre-DIONE.

In conclusion, TRANsCre-DIONE proves to be highly efficient and specific in transdifferentiating reactive astrocytes to appropriate neurons in different regions of the injured brain. The complete absence of tumorigenesis and no need for transplantation are the biggest advantages of TRANsCre-DIONE over stem-cell approaches. Future improvements should include combining the three components of TRANsCre-DIONE into one AAV vector for easy handling and reduced cost of virus production. Finally, the feasibility of TRANsCre-DIONE in the monkey brain promises immediate clinical applications in various traumatic brain injuries and neurodegenerative diseases to replenish dead neurons and regenerate damaged brain.

## Supporting information

Supplemental Informations

## ACKNOWLEDGMENTS

This research was supported by Institute for Basic Science (IBS) (IBS-R001-D2), and Creative Research Initiative Program, National Research Foundation (NRF) of Korea (2015R1A3A2066619).

## AUTHOR CONTRIBUTIONS

H.Y.A., H-L.L, Y.H. and C.J.L. designed the study. H.Y.A and H-L.L. analyzed the data. H.Y.A. and H-L.L. and C.J.L wrote the manuscript. H.Y.A. and J.P.H. cloned all the viral vectors. H.Y.A. performed all surgery, IHC, analysis with mouse brain part. H.Y.A., D.W.C., Y.S.Y. and S.C.H. performed and supported the animal surgery, caring, sacrificing with monkey. H.Y.L. and H-L.L. performed all animal surgery behavioral tests, and analysis with SCI mouse model. H.Y.A., J.M.L., J.S.W. and J.K.L. performed slice electrophysiology. M.G.P. designed the schematic figures. All authors contributed to discussion of the results.

## METHOD & METARIALS

### Animals

To differentiate of reactive astrocytes into neurons directly in the brain, 7 to 10-week-old C57BL/6J mice (DBL; Daehan Bio Link, Korea) were used for immunohistochemistry and electrophysiology experiments. Mice were housed in an animal facility permitted by the Association for Assessment and Accreditation of Laboratory Animal Care (AAALAC). All animals were maintained in a vivarium with light/dark cycle (8:00 AM∼8:00 PM). Animal care and handling were performed according to the directives of the Animal Care and Use Committee and institutional guidelines of KIST (Seoul, Korea). Animals were randomly used for experiments.

Three- to 4-year old cynomolgus monkeys (*Macaca fascicularis*, 2 males) were purchased from NafoVanny (Dong Nai Province, Vietnam), and they were 4.1 and 3.9 kg on the dosing day. The monkeys were housed individually in a stainless-steel cage (543 mm width x 715 mm depth x 818 mm height) and acclimatized in the study room after a 30-day quarantine period. Throughout the study period, supplements, toys, and stainless-steel mirrors were supplied for environmental enrichment. The room conditions were automatically controlled based on standard operating procedures (a temperature of 20 – 29 °C, relative humidity of 30 – 70 %, a lighting cycle of 12 hours dark/12 hours light at 300 – 700 lux and 10 – 20 air changes per hour). Filtered, ultraviolet light-irradiated municipal tap water was allowed ad libitum and 120 g of commercial monkey chow (Certified Primate Diet #5048, PMI Nutrition International, USA) was provided daily. The animals were identified using body tattoos and cage cards. This study was performed in facilities approved by the Association for Assessment and Accreditation of Laboratory Animal Care International. All procedures were approved by the Institutional Animal Care and Use Committee of the Korea Institute of Toxicology.

### DNA Cloning and virus packaging

cDNAs encoding for full-length *Lcn2* (GenBank accession no. NM_008491.1) and *Neurog2* (NM_008430) were obtained by using an RT–PCR based cloning from mouse brain. *iNOS* promoter (NM_000625, Product ID: HPRM30585) was purchased from Genecopoeia. GFAP-Ccre, GFAP-Ncre vectors were kindly provided by Dr. Rolf Sprengel (Max Planck Institute for Medical Research). pAAV-GFAP::Ccre, pAAV-GFAP::Ncre, pAAV-Lcn2::Ncre, pAAV-iNOS::Ncre, pAAV-EF1α::DIO-Neurog2-IRES-GFP, pAAV-EF1α::DIO-GFP were packaged into AAV.

### Astrocytes primary culture

Primary cultured astrocytes were electroporated at 4 DIV with a mixture of 3 plasmids (pAAV-EF1α-df-Ngn2-IRES-GFP, pAAV-Lcn2-Ncre, pAAV-GFAP-Ccre, 1:1:1, 5μg each), astrocyte media was changed to neuronal media 8 days after gene transfection. Astrocytes media contains 10% HI Horse serum (Gibco, 26050-088), 10% HI Fetal Bovine Serum (Gibco, 10082-147), 1% Penicillin-Streptomycin (Gibco, 15140-122) in Dulbecco Modified Eagle Medium (DMEM, Corning, 10-013-CVR), neuronal media inducing neuronal differentiation includes 2% B27 (Invitrogen, 17504-044), 2% Glutamax (Gibco, 35050-061) in F-12 media (Gibco, 11765-054).

### Stereotaxic surgery in Brain

#### Mouse

7-10 -week-old 57/BL6 mice were used. Mice (8–10 weeks old) were anesthetized by intraperitoneal injection of 2% avertin (20 μl g − 1) and placed into stereotaxic frames. AAV (titer: 1.5×10^12^ GC) was loaded into a 22G Hamilton syringe and injected bilaterally into the mouse striatum region (−0.5 mm AP, ± 1.85 mm ML, −3.0 mm DV from the dura) at a rate of 0.2 μl min − 1 (total 2 μl) with a 25 μl syringe using a syringe pump (KD Scientific, Holliston, MA, USA).

#### Monkey

Animals (age 2-6 years) were anesthetized with approximately 8 – 10 mg/kg of zoletil 50 (Virbac Korea) 30 min before clamped to the stereotaxic instrument (David Kopf instrument, model 1404). Total 25 uL virus was injected using the Hamilton syringe and infusion pump at a speed of 2.5 uL per minutes. After surgery, monkeys were prescribed 20 mg/kg of Cephalosporin-C twice per day for 5 days, 2 mg/kg of Ketoprofen once a day for 3 days and 1 mg/kg of Prednisolone acetate once a day for 5 days. To prevent infection following surgery, surgical sites were disinfected with Betadine for 5 days. A mixture of AAV (2/5)-iNOS-Ccre, AAV (2/5)-GFAP-Ncre and AAV (2/5)-DF-GFP and a mixture of AAV (2/5)-iNOS-Ccre, AAV (2/5)-GFAP-Ncre and AAV (2/5)-DF-NGN2-IRES-GFP were injected in the putamen (AP +0.9, ML +09-1.1, DV-2.7 from bregma).

### Establishment of Spinal cord injury (SCI) model and AAV injection

For the establishment of SCI model, *C57BL/6* was used (20g ± 2g; OrientBio, Kyungki-do, Korea), housed in an animal facility permitted by the Association for Assessment and Accreditation of Laboratory Animal Care (AAALAC). All animals were anesthetized with ketamine (100mg/kg; Yuhan, Seoul, Republic of Korea), Rompun (10mg/kg; Bayer Korea, Seoul, Republic of Korea) and isotropy 100 (1 to 3%, Troikaa Pharmaceuticals Ltd, Gujarat, India). Laminectomy was performed at thoracic number 10 to expose the spinal cord. The spinal cord was compressed for 3s by self-closing forceps (Fine Science Tools, North Vancouver, Canada). After spinal cord injury, muscle and skin were closed using 6/0 non-absorbable nylon sutures. Sham animals were operated the only laminectomy. Spinal cord injured animals were randomly separated to PBS, Ctrl, and Ngn2. At 2 weeks after injury, 1 μl PBS and AAV were injected at two locations (1 mm proximal and distal to the injured site) using by 33gauge Hamilton syringe. The injection rate was constantly maintained as 0.2 μl to min. After injection, the needle was kept at the injection site for additional 5 min.

### BrdU assay

For detection of cell proliferation, BrdU (50mg/kg; Sigma-Aldrich, B5002) was administrated by intraperitoneal injection. BrdU was injected daily for 7 days from 1 week and 7 weeks after injury. BrdU injected mice were sacrificed at the last day of BrdU injection. The concentration of BrdU solution was 10 mg/ml in 0.9% saline and filtrated before injected to the mouse. BrdU was detected by immunohistochemistry using the mouse anti-BrdU antibody (Thermo Fisher, ZBU30; 1:2000).

### Behavioral test

For confirmation of locomotor recovery, the behavioral test was performed every week from 1 week to 7 weeks after injury. The Basso Mouse Scale (BMS) score was used to quantify of hind limb movement during open field locomotion. The BMS score is separated as 0 to 9 scores. The 0 score means complete paralysis and 9 score means normal locomotion.

### Tissue sample preparation for Eriochrome Cyanine (EC) staining and Immunohistochemistry

For isolation of spinal cord, animals were sacrificed at 3 weeks and 8 weeks after injury. Blood was completely removed by saline perfusion and fixed by 4% paraformaldehyde (PFA, Duksan, Ansan, Korea). The isolated tissues were fixed in 4% PFA for overnight and transferred to 30% Sucrose for dehydration at 4°C. Dehydrated tissues were embedded in optimal cutting temperature compound (Sakura Finetek, Torrance, CA, USA), frozen at −80°C, and cut into a thickness of 20 Lm by cryostat.

### Eriochrome Cyanine (EC) staining

The sectioned tissues were dried on room temperature (RT) for 2 h and immersed in acetone for 5 min. After immersion, tissues were incubated at RT for 10 min and stained by EC solution (MERCK, Kenilworth, NJ, USA) at RT for 30 min. The stained tissues were rinsed by running water and 5% iron alum (Sigma-Aldrich, F3629) until observing the gray matter. Tissues were differentiated by borax-ferricyanide solution (Borax, Sigma-Aldrich, 71997; Potassium ferricyanide, Sigma-Aldrich, 702587) and dehydrated by graded ethanol solutions of 70%, 90%, and 100%. After all process, tissues were mounted by permanent mounting medium (Fisher Scientific, Hampton, NH, USA) and observed by the light microscope (DM 2500, Leica, Wetzlar, Germany).

### Immunohistochemistry

#### Immunohistochemistry in Brain

Animals were euthanized upon completion of the treatment period with an overdose of thiopental sodium administered intravenously the bled by blood collection via posterior vena cava. They were fasted over 16 hours before the necropsy. The animals were examined carefully for external abnormalities. The abdominal, thoracic and cranial cavities were examined for abnormalities and the brain then removed and examined. Adult mice were deeply anesthetized with 2% avertin (20 µg/g) and perfused with 0.1M PBS (Phosphate buffered saline) followed by ice cold 4 % PFA (paraformaldehyde). Excised brains were post-fixed overnight in 4 % PFA at 4 °C and immersed in 30% sucrose for 24 hrs for cryo-protection. The 30-50 µm coronal sections were prepared with cryo-stat microtome (HM525, Thermo Fisher Scientific). Sections with were rinsed in PBS three times and incubated 1 hr at RT with blocking solution (0.3% Triton-X, 2 % normal serum in 0.1 M PBS). Sections were incubated overnight in a mixture of the primary antibodies with blocking solution at 4 °C on shaker. After washing three times in PBS, sections were incubated with corresponding secondary antibodies for two hours and then rinsed three times with PBS. If needed, DAPI staining was added during second wash step. (Pierce, 1:3000) After being dried, stained tissues were mounted with an anti-fade mounting medium (Dako). A series of fluorescence images were obtained with a confocal microscope (Nikon, A1R) and images were processed for later analysis using ImageJ program imaging software.

#### Immunohistochemistry in Spinal Cord

The tissue sections were washed three times by 0.3% Tween 20 (Sigma-Aldrich, P1379) in PBS at RT for 5 min each and blocked by 10% normal donkey serum (Jackson ImmunoResearch, West Grove, PA) in PBS containing 0.3% Triton X-100 (Sigma-Aldrich, X100) at RT for 1 h. After then, primary antibodies were treated such as goat anti-GFP (Abcam, Cambridge, UK, ab6662, 1:1000), chicken anti-GFAP (Abcam, ab4674, 1:2000), chicken anti-MBP (Abcam, ab134018, 1:400), rabbit anti-MAP2 (Abcam, ab32454, 1:1000), rabbit anti-NeuN (Abcam, ab177487, 1:800) and mouse anti-islet1(DSHB Iowa City, IA, USA, 39.4D5, 1:100) and incubated at 4°C for overnight. After three times 0.3% Tween 20 washing, sections were incubated with species-specific secondary antibodies conjugated with fluorescent agents such as Cy™3-donkey anti-mouse IgG (H+L) (Jackson ImmunoResearch, 175-165-151; 1:600), Cy™3-donkey anti-chicken IgG (H+L) (Jackson ImmunoResearch, 175-165-155; 1:600), Alexa Fluor 647-donkey anti-rabbit IgG (H+L) (Jackson ImmunoResearch, 711-606-152; 1:300), Cy™5-donkey anti-chicken IgG (H+L) (Jackson ImmunoResearch, 703-175-155; 1:300) and DyLight™405-donkey anti-chicken IgY_++_ (H+L) (Jackson ImmunoResearch, 703-476-155; 1:600) at RT for 1 h. incubated sections were washed three times by 0.3% Tween 20 and stained with DAPI (DAPI; Vector Laboratories, Peterborough, UK). The fluorescence was observed under a confocal laser scanning microscopy (LSM700, Carl Zeiss, Oberkochen, Germany).

### Electrophysiology

#### Slice preparation

Coronal mouse brain slices (300 μm) containing trigeminal caudal nucleus region were acutely prepared from C57/BL6 (age 7∼10 weeks). Following decapitation the brain was rapidly removed and placed in cold artificial cerebrospinal fluid (ACSF) having the following composition (in mM): 130 NaCl, 24 NaHCO3, 3.5 KCl, 1.25 NaH2PO4, 1 CaCl2, 3 MgCl2 and 10 glucose, pH 7.4; room temperature with oxygenation (95% O2 and 5% CO2). The slices were made using an oscillating tissue slicer (DSK LinearSlicer, Kyoto, Japan) at 4°C and stored in room temperature with oxygenation (95% O2, 5% CO2) and prepared slices were left to recover for at least 1 hour before recording.

#### Mouse brain ex-vivo

Each slice that was studied was transferred from a recovery/holding reservoir to the recording chamber of a fixed-stage upright microscope (<u>Olympus</u>) and submerged in oxygenated ACSF that was supplied to the chamber at a rate of 1.5∼2 ml/min. The submerged slice was visualized either directly via the microscope’s optics, or indirectly via a high-resolution CCD camera system (Orca Flash 2.1, Hamamatsu) that received the output of a CCD camera attached to the microscope’s video port. Experiments with a holding current of more than −100 pA or in which there was a change in input resistance >30% of the control were rejected. Recordings were obtained using Multiclamp 700A amplifier (Molecular Devices, Sunnyvale, CA) and were filtered at 2 kHz. Current recordings under ramp protocol and step were digitized at 10 kHz with DigiDATA 1550B1 (Molecular Devices, Sunnyvale, CA) and analyzed using pCLAMP 10 software (Molecular Devices, Sunnyvale, CA). Whole-cell recordings from GFP expressing cells were carried out with internal solution composed of (mM): 120 potassium gluconate, 10 KCl, 1 MgCl2, 0.5 EGTA, 40 HEPES (pH 7.2 was adjusted with KOH)

## Statistics

Data are presented as mean ± SEM. Levels of statistical significance are indicated as follows: *P < 0.05, **P < 0.01, and ***P < 0.001. For full information, see statistics tables in supplementary information.

### KEY RESOURCES TABLE

| REAGENT or RESOURCE | SOURCE | IDENTIFIER |
| --- | --- | --- |
| **Antibodies** |  |  |
| Chicken anti-GFAP antibody | Millipore | AB5541 |
| Chicken anti-MAP2 antibody | abCam | ab5392 |
| Mouse anti-NeuN antibody | Millipore | MAB377 |
| Rabbit anti-iNOS antibody | Thermo Fisher | PA3-030A |
| Goat anti-Lcn2 antibody | abCam | ab2267 |
| Rabbit anti-islet1 antibody | DSHB Iowa City | N/A |
| Chicken anti-MBP antibody | abCam | ab134018 |
| Goat anti-GFP | abCam | ab6662 |
| Mouse anti-BrdU antibody | Thermo Fisher | ZBU30 |
| **Bacterial and Virus Strains** |  |  |
| AAV-Lcn2-Ncre | KIST virus facility | N/A |
| AAV-GFAP-Ccre | KIST virus facility | N/A |
| AAV-iNOS-Ccre | KIST virus facility | N/A |
| AAV-GFAP-Ncre | KIST virus facility | N/A |
| AAV-EF1α-df-Ngn2-IRES-EGFP | KIST virus facility | N/A |
| AAV-EF1α-df-GFP | KIST virus facility | N/A |
| **Chemicals, Peptides, and Recombinant Proteins** |  |  |
| Dulbecco Modified Eagle Medium (DMEM) | Corning | 10-013-CVR |
| Penicillin-Streptomycin | Gibco | 15140-122 |
| HI Fetal Bovine Serum | Gibco | 10082-147 |
| HI Horse serum | Gibco | 26050-088 |
| B27 | Invitrogen | 17504-044 |
| Glutamax | Gibco | 35050-061 |
| F-12 media | Gibco | 11765-054 |
| BrdU | Sigma-Aldrich | B5002 |
| Iron alum | Sigma-Aldrich | F3629 |
| Borax | Sigma-Aldrich | 71997 |
| Potassium ferricyanide | Sigma-Aldrich | 702587 |
| **Recombinant DNA** |  |  |
| pAAV-Lcn2-Ncre |  |  |
| pAAV-GFAP-Ccre |  |  |
| pAAV-iNOS-Ccre |  |  |
| pAAV-GFAP-Ncre |  |  |
| pAAV-EF1α-df-Ngn2-IRES-EGFP | | |
| pAAV-EF1α-df-GFP | | |
| **Software and Algorithms** |  |  |
| GraphPad Prism 7 | GraphPad software | https://www.graphpad.com/scientific-software/prism/;<br>RRID: SCR_015807 |
| ImageJ | National Institutes of Health | https://imagej.nih.gov/ij/; RRID: SCR_003070 |
| **Other** |  |  |
| Neon Transfection System |  | MPK5000 |
| Neon™ Transfection System Kit |  | MPK10025 |

