## Supplemental Figures for "TRANsCre-DIONE transdifferentiates scar-forming reactive astrocytes into functional motor neurons"

### Figure S1

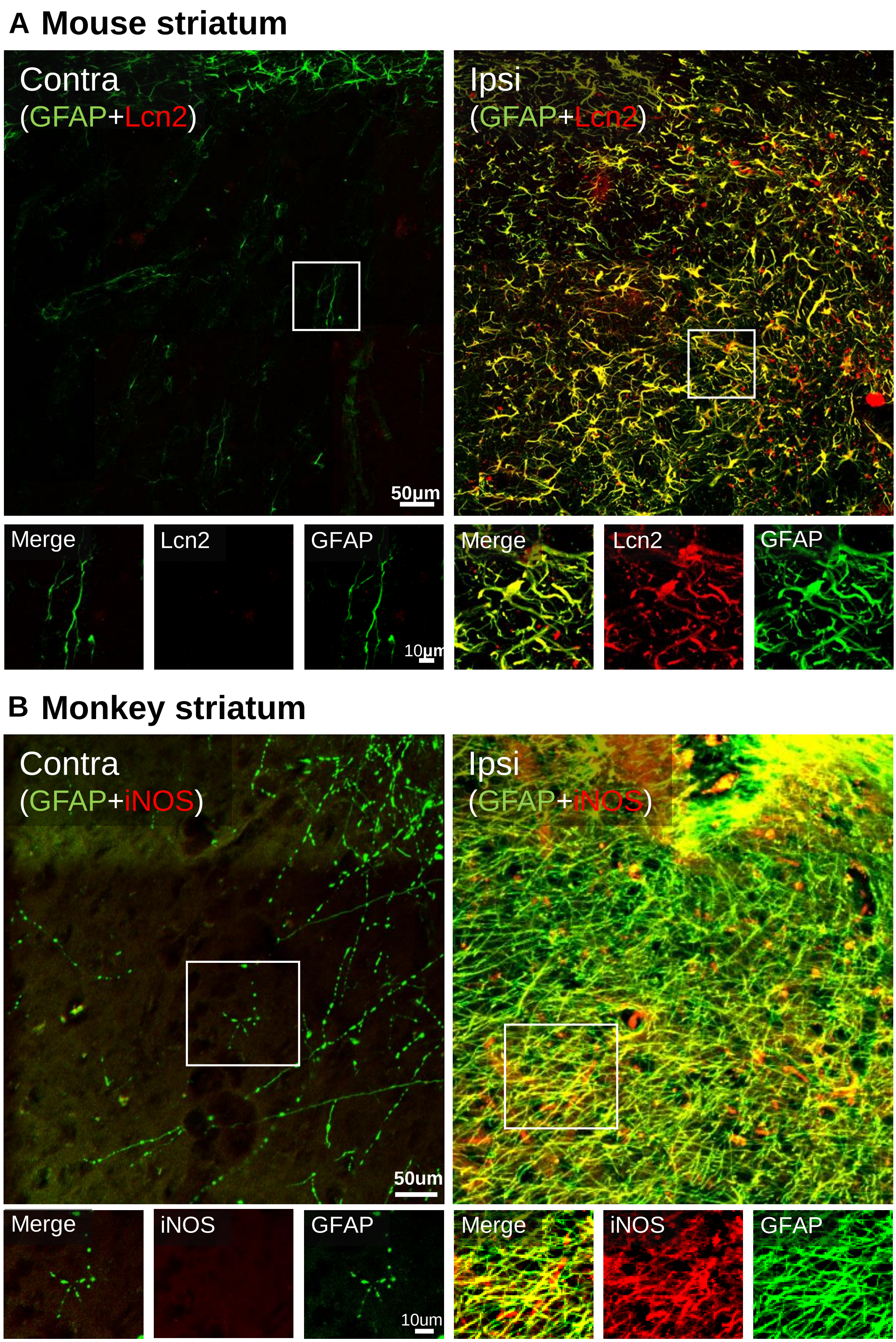


**Figure S1. Lcn2 or iNOS with GFAP expression in contra or ipsi lateral hemispheres, related with figure 1**

(A) Colocalization of Lcn2 (red) and GFAP (green) in contra lateral (left) and ipsi lateral (right) striatum of mouse. Lower panel shows magnified images. Scale bar=50 μm, 10 μm (magnified images) (B) Colocalization of iNOS (red) and GFAP (green) in contra lateral (left) and ipsi lateral (right) putamen of cynomolgus monkey. Lower panel shows magnified images. Scale bar=50 μm, 10 μm (magnified images).

### Figure S2

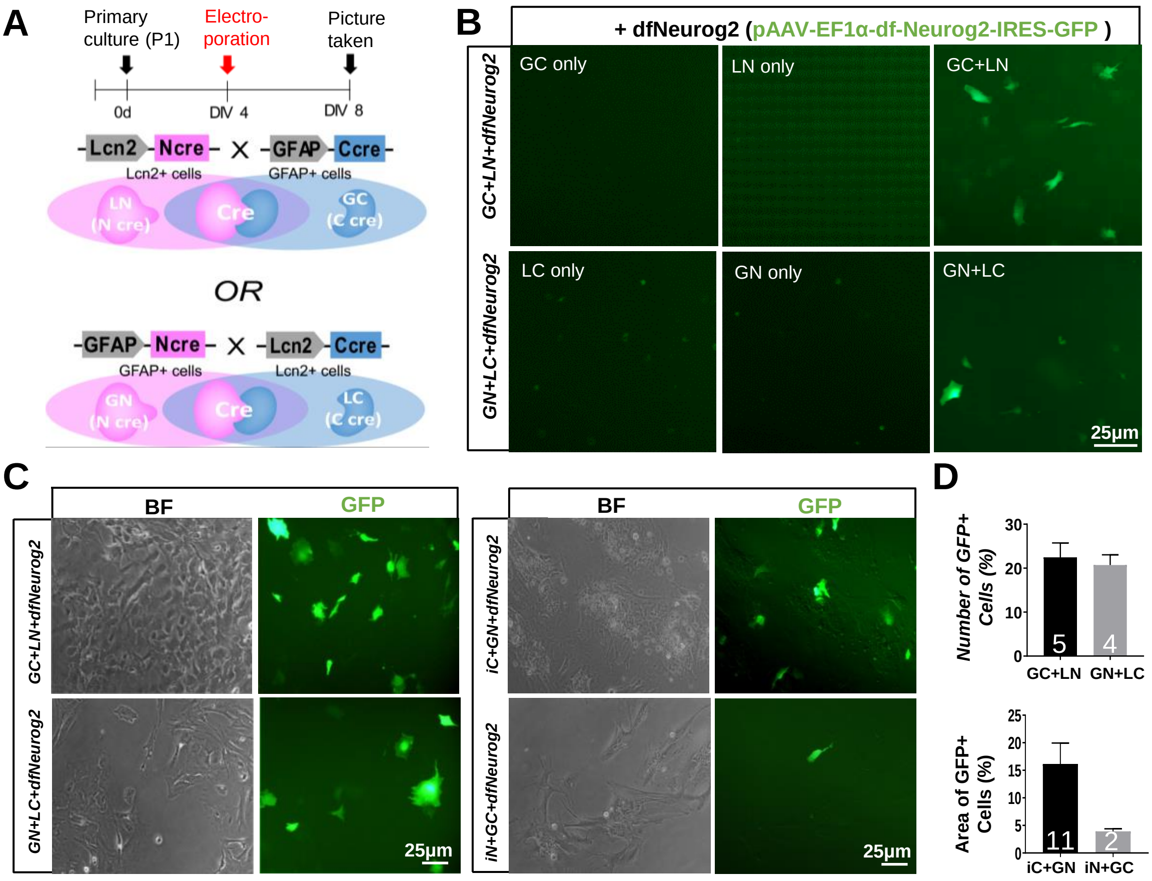


**Figure S2. Cell-specificity and expression-efficiency of Split-Cre, related with figure 1**

(A) Experimental protocol for working test of the split-Cre system in primary cultured astrocytes. Electroporation was performed DIV4 and the fluorescence was detected DIV 8 (4 days after electroporation). (B) pAAV-EF1α-df-Ngn2-IRES-GFP (dfNgn2; GFP) was electroporated with Ccre (GC, GFAP-Ccre; LC, Lcn2-Ccre) or Ncre (LN, Lcn2-Ncre; GN, GFAP-Ncre) only or both. Scale bar= 25 μm. (C) Efficiency test for expression of each pair of split-Cre with dfNgn2. GFAP and Lcn2 promoters, the pair of GC+LN and GN+CL (left). iNOS and GFAP promoters, the pair of iC+GN and iN+GC (right). Bright field (BF) images show cell density, GFP fluorescence (GFP) shows the gene expression. Scale bar=25 μm. (D) Summary bar graphs showing the percentage of the GFP-expressing cells or area in the fields of views. n= 5 (GC+LN), 4 (GN+LC). n= 11 (iC+GN), 2 (iN+GC).

### Figure S3

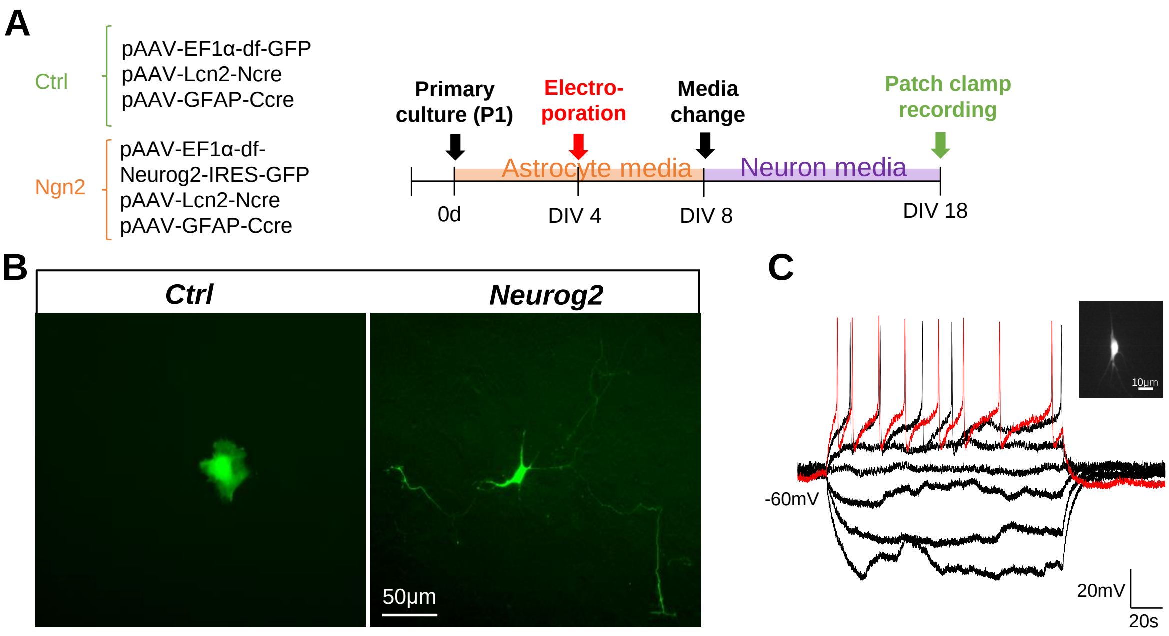


**Figure S3. Transdifferentiation of reactive astrocytes into functional neurons in culture, related with figure 2**

(A) A mixture of 3 vectors (TRANsCre-DIONE, 5 μg each) were electroporated in primary cultured astrocytes for transdifferentiation of reactive astrocytes into neurons and the timeline of transfection and treating neuronal media. Electroporation was performed at days *in vitro* (DIV) 4, media was changed to neuronal media 4 days after electroporation in Neurog2 group. The transdifferentiated cells were recorded by whole-cell patch-clamp recording at DIV 18. (B) Transdifferentiated cells (GFP) show the neuron-like morphology 14 days after electroporation (DIV18) in Neurog2 group. Scale bar= 50μm. (C) Whole-cell patch clamp recording with an GFP expressing cell (inset, scale bar= 10μm) was performed at DIV 18. Neuron-like cell shows action potentials, but not spontaneous. Scale bar= 20 mV, 20 sec.

### Figure S4

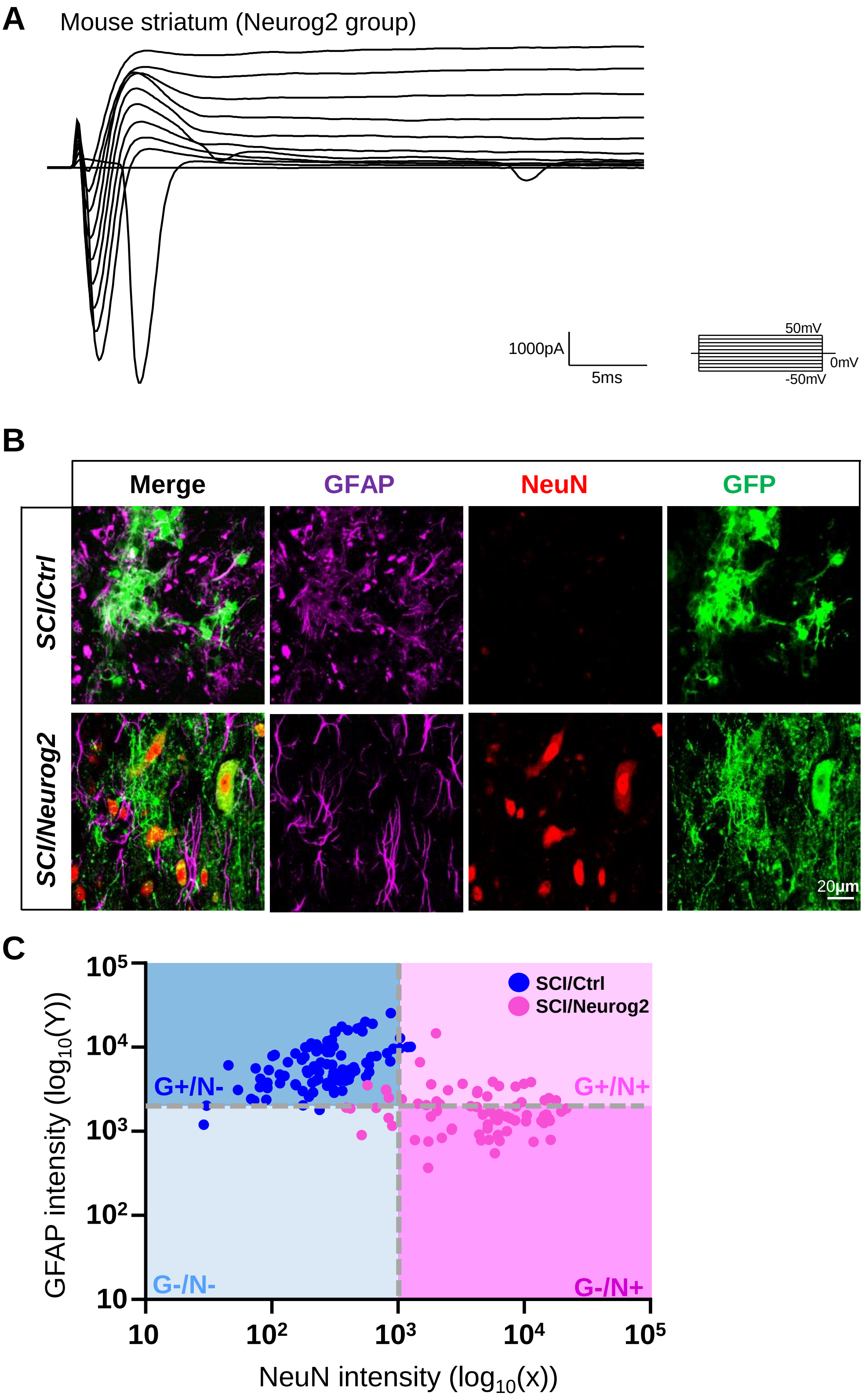


**Figure S4. Transdifferentiated functional neurons in mouse striatum and the efficiency of transdifferentiation in SCI, related with figure2 and figure 6**

(A) Na-K currents were recorded by voltage-clamp step protocol in GFP-expressing cells in mouse striatum. (B) Colocalization of GFAP (magenta), NeuN (red) and GFP (green) in SCI/Ctrl and SCI/Neurog2 groups. Scale bar= 20 μm. (C) Scatter plot of GFP-expressing cells classified by GFAP and NeuN intensity from SCI/Ctrl (blue,) and SCI/Neurog2 (pink,) groups. GFP+ cells were divided in four groups, G+/N- (blue), G-/N- (light blue), G+/N+ (light pink), G-/N+ (pink). n=102 (SCI/Ctrl), 77 (SCI/Neurog2).

### Figure S5

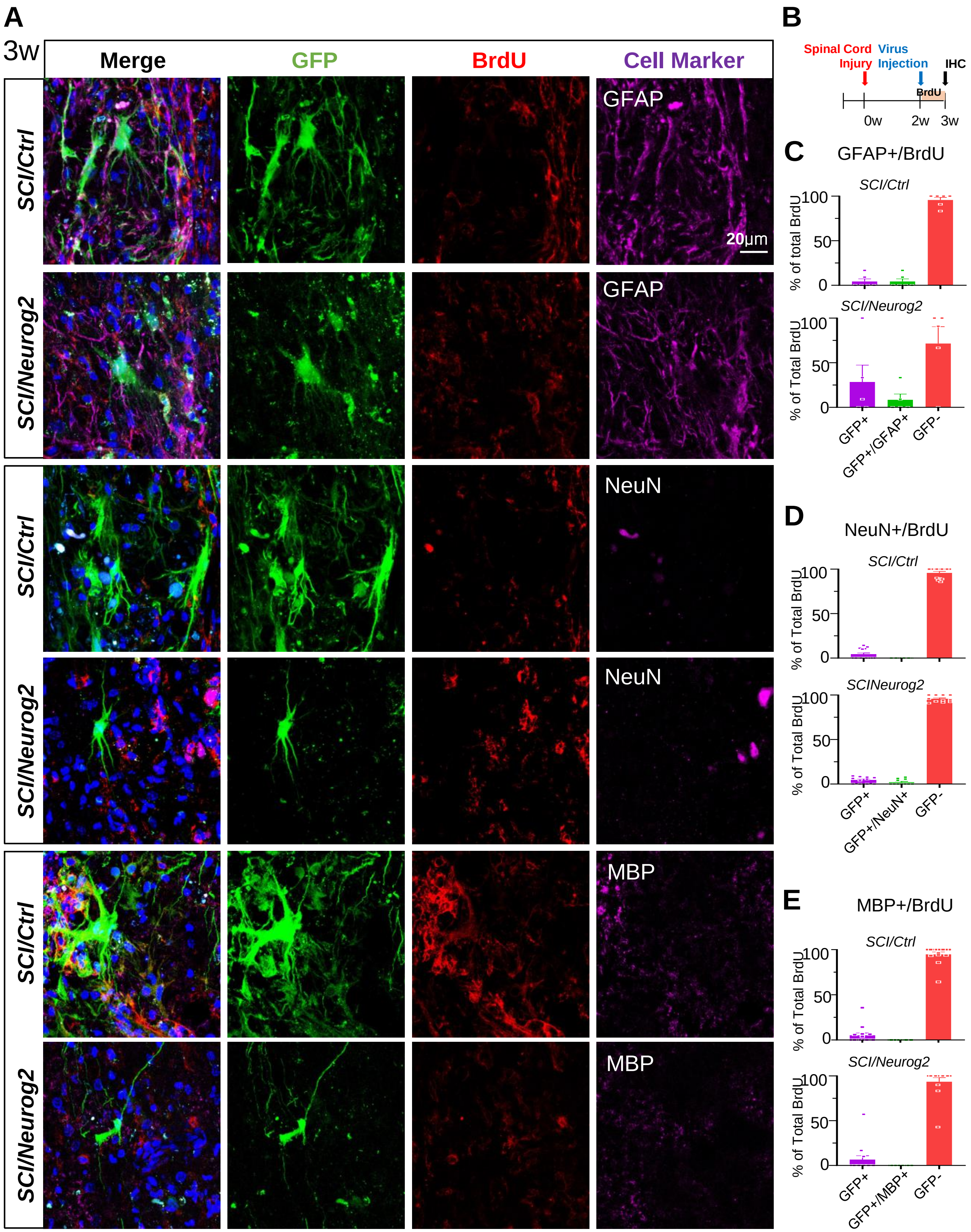


**Figure S5. Transdifferentiated neurons are mostly originated from non-proliferating cells, related with figure 7**

(A) Colocalization of BrdU (red, treated for 1week from 2-3 weeks after injury) in GFP-expressing cells (green) with cell markers (magenta; GFAP, upper; NeuN, middle; MBP, lower) in SCI/Ctrl and SCI/Neurog2 groups. The mice were sacrifice 3 weeks after injury (3w). Scale bar=20 μm. (B) Timeline of treating BrdU and IHC after sacrificing in SCI mice. (C) Scattered bar graph showing comparison of percentage of BrdU-positive cells in GFP+, GFAP+/GFP+ and GFP- cells in SCI/Ctrl and SCI/Neurog2 groups. n=7 (SCI/Ctrl group), 17 (SCI/Neurog2 group). (D) Scattered bar graph showing comparison of percentage of BrdU-positive cells in GFP+, NeuN+/GFP+ and GFP- cells in SCI/Ctrl and SCI/Neurog2 groups. n=7 (SCI/Ctrl group), 17 (SCI/Neurog2 group). (E) Scattered bar graph showing comparison of percentage of BrdU-positive cells in GFP+, MBP+/GFP+ and GFP- cells in SCI/Ctrl and SCI/Neurog2 groups. n=7 (SCI/Ctrl group), 17 (SCI/Neurog2 group).

### Figure S6

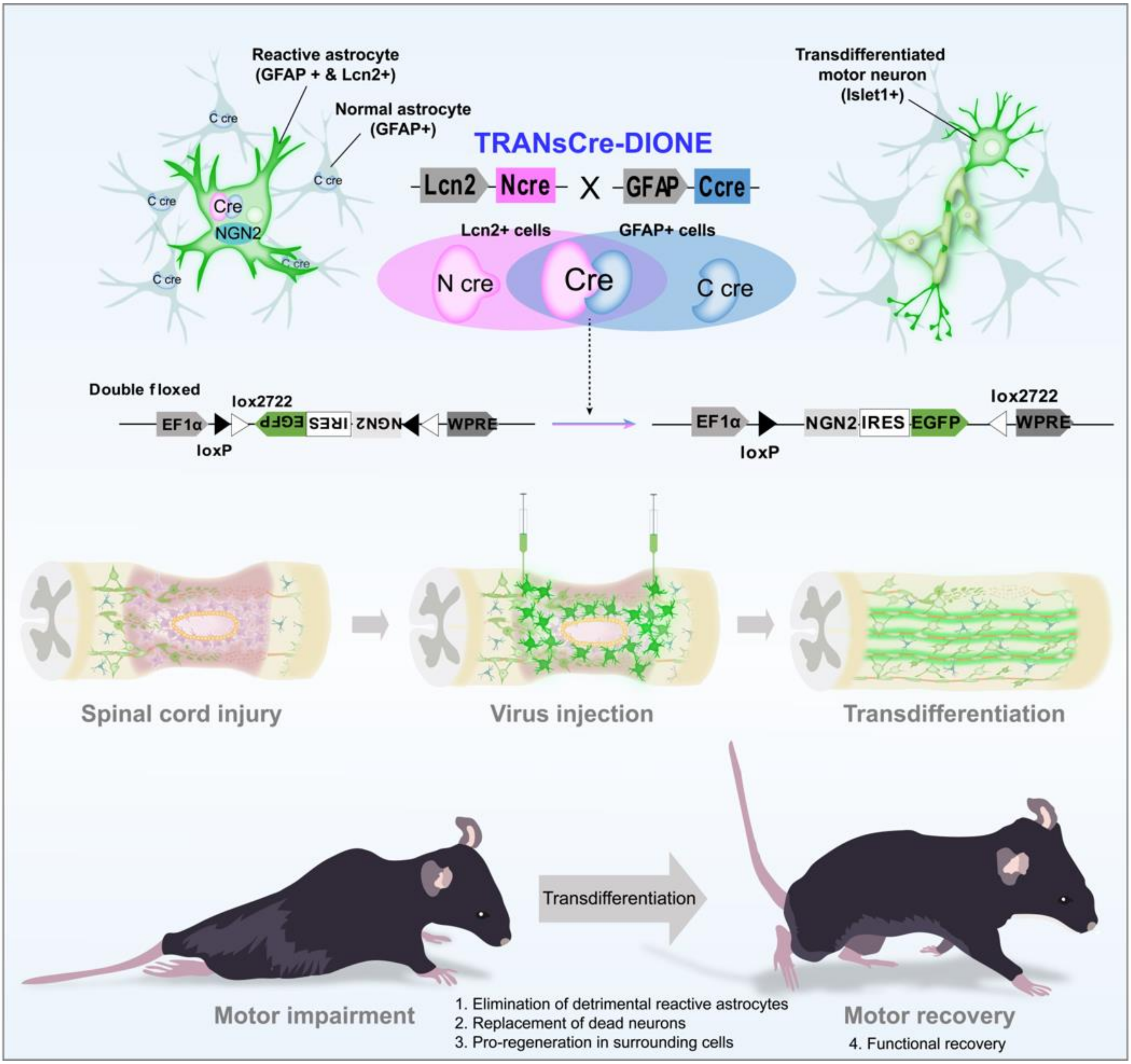


**Figure S6. Schematic principle of treating SCI via direct reprogramming of reactive astrocytes into motor neurons.**
