## Supplemental Informations for "TRANsCre-DIONE transdifferentiates scar-forming reactive astrocytes into functional motor neurons"

Table S1

Dunnett's multiple comparison test, with individual variances computed for each comparison

Mixed-effect model with the Geisser-Greenhouse correction, matched values are spread across a row

|  |  | **Group** | **Number (%)** | |
| --- | --- | --- | --- | --- |
|  |  |  | **Ctrl** | **Neurog2** |
| Fig.  2.C. | Number of cells | NeuN | 1 (1.07) | 12 (8.89) |
|  |  | GFAP | 83  (89.247) | 39 (28.89) |
|  |  | Undefined | 9 (9.677) | 84 (62.22) |
|  |  | **In NeuN+/GFP+** | | |
|  |  | **Group** | **GABA+** | **GABA-** |
| Fig.  2.E. | Number of cells | Average (%) | 29.33 | 14.16 |
|  |  | SEM | 71.33 | 125 |

Table S2

Dunnett's multiple comparison test, with individual variances computed for each comparison

Mixed-effect model with the Geisser-Greenhouse correction, matched values are spread across a row

|  |  | **Group** | **Number** | **Intensity of GFP+ cells (Mean±SEM)** | **t-test**  **(P value)** |
| --- | --- | --- | --- | --- | --- |
| Fig.3.E | Ctrl | NeuN+ | 2 | 488.4987±29.0234 | <0.0001 |
|  |  | GFAP+ | 96 | 2337.581±93.8979 |  |
| Fig.3.F | Neurog2 | NeuN+ | 99 | 1040.552±45.4978 | <0.0001 |
|  |  | GFAP+ | 125 | 1555.643±75.1399 |  |
| Fig.3.G | NeuN+ | Ctrl | 2 | 488.4987±29.0234 | <0.0001 |
|  |  | Neurog2 | 99 | 1040.552±45.4978 |  |
| Fig.3H | GFAP+ | Ctrl | 96 | 2337.581±93.8979 | <0.0001 |
|  |  | Neurog2 | 125 | 1555.643±75.1399 |  |

Table S3

Dunnett's multiple comparison test, with individual variances computed for each comparison

Mixed-effect model with the Geisser-Greenhouse correction, matched values are spread across a row

| Fig.  4.C. |  | **Group** | **Number** | **Mean±SEM** | **Multiple Comparisons** | ***P* value** |
| --- | --- | --- | --- | --- | --- | --- |
|  | 1 Week | Sham | 5 | 9±0.000 | SCI/Neurog2 vs. Sham | <0.0001 |
|  |  | SCI/PBS | 10 | 0.175±0.124 | SCI/Neurog2 vs. SCI/PBS | 0.8054 |
|  |  | SCI/Ctrl | 10 | 0.275±0.142 | SCI/Neurog2 vs. SCI/Ctrl | 0.9936 |
|  |  | SCI/Neurog2 | 11 | 0.3182±0.143 |  |  |
|  | 2 Week | Sham | 5 | 9±0.000 | SCI/Neurog2 vs. Sham | <0.0001 |
|  |  | SCI/PBS | 10 | 0.475±0.126 | SCI/Neurog2 vs. SCI/PBS | 0.7225 |
|  |  | SCI/Ctrl | 10 | 0.45±0.082 | SCI/Neurog2 vs. SCI/Ctrl | 0.4356 |
|  |  | SCI/Neurog2 | 11 | 0.6136±0.091 |  |  |
|  | 3 Week | Sham | 5 | 9±0.000 | SCI/Neurog2 vs. Sham | <0.0001 |
|  |  | SCI/PBS | 10 | 0.925±0.261 | SCI/Neurog2 vs. SCI/PBS | 0.0346 |
|  |  | SCI/Ctrl | 10 | 0.6±0.107 | SCI/Neurog2 vs. SCI/Ctrl | 0.0002 |
|  |  | SCI/Neurog2 | 11 | 1.841±0.203 |  |  |
|  | 4 Week | Sham | 5 | 9±0.000 | SCI/Neurog2 vs. Sham | <0.0001 |
|  |  | SCI/PBS | 10 | 1.075±0.253 | SCI/Neurog2 vs. SCI/PBS | 0.0101 |
|  |  | SCI/Ctrl | 10 | 1.025±0.146 | SCI/Neurog2 vs. SCI/Ctrl | 0.0020 |
|  |  | SCI/Neurog2 | 11 | 2.273±0.258 |  |  |
|  | 5 Week | Sham | 5 | 9±0.000 | SCI/Neurog2 vs. Sham | <0.0001 |
|  |  | SCI/PBS | 10 | 0.9±0.155 | SCI/Neurog2 vs. SCI/PBS | <0.0001 |
|  |  | SCI/Ctrl | 10 | 1.3±0.128 | SCI/Neurog2 vs. SCI/Ctrl | 0.0008 |
|  |  | SCI/Neurog2 | 11 | 2.523±0.230 |  |  |
|  | 6 Week | Sham | 5 | 9±0.000 | SCI/Neurog2 vs. Sham | <0.0001 |
|  |  | SCI/PBS | 10 | 1.1±0.150 | SCI/Neurog2 vs. SCI/PBS | <0.0001 |
|  |  | SCI/Ctrl | 10 | 1.15±0.205 | SCI/Neurog2 vs. SCI/Ctrl | 0.0002 |
|  |  | SCI/Neurog2 | 11 | 2.773±0.249 |  |  |
|  | 7 Week | Sham | 5 | 9±0.000 | SCI/Neurog2 vs. Sham | <0.0001 |
|  |  | SCI/PBS | 10 | 1.125±0.198 | SCI/Neurog2 vs. SCI/PBS | <0.0001 |
|  |  | SCI/Ctrl | 10 | 1.325±0.149 | SCI/Neurog2 vs. SCI/Ctrl | <0.0001 |
|  |  | SCI/Neurog2 | 11 | 3.136±0.114 |  |  |
|  | 8 Week | Sham | 5 | 9±0.000 | SCI/Neurog2 vs. Sham | <0.0001 |
|  |  | SCI/PBS | 10 | 1.225±0.188 | SCI/Neurog2 vs. SCI/PBS | <0.0001 |
|  |  | SCI/Ctrl | 10 | 1.325±0.149 | SCI/Neurog2 vs. SCI/Ctrl | <0.0001 |
|  |  | SCI/Neurog2 | 11 | 3.182±0.169 |  |  |

Table S4

Multiple comparison for unpaired non-parametric one-way ANOVA with Kruskal-Walis test

|  |  | **Group** | **Number** | **Mean Area (±SEM, Normalized by Sham)** | **Multiple Comparisons** | ***P* value** |
| --- | --- | --- | --- | --- | --- | --- |
| Fig.  5.B. | Gray matter area | Sham | 10 | 1±0.03390 | SCI/Neurog2 vs. Sham | 0.0725 |
|  |  | SCI/PBS | 8 | 0.2798±0.08361 | SCI/Neurog2 vs. SCI/PBS | 0.0446 |
|  |  | SCI/Ctrl | 19 | 0.3774±0.03225 | SCI/Neurog2 vs. SCI/Ctrl | 0.1721 |
|  |  | SCI/Ngn2 | 13 | 0.6637±0.1105 |  |  |
|  | White matter area | Sham | 10 | 1±0.05280 | SCI/Neurog2 vs. Sham | 0.0443 |
|  |  | SCI/PBS | 8 | 0.2877±0.03132 | SCI/Neurog2 vs. SCI/PBS | 0.6995 |
|  |  | SCI/Ctrl | 18 | 0.1270±0.03513 | SCI/Neurog2 vs. SCI/Ctrl | 0.0019 |
|  |  | SCI/Ngn2 | 13 | 0.4952±0.05925 |  |  |
|  | Total area | Sham | 10 | 0.9670±0.02681 | SCI/Neurog2 vs. Sham | 0.0464 |
|  |  | SCI/PBS | 8 | 0.2840±0.04075 | SCI/Neurog2 vs. SCI/PBS | 0.3312 |
|  |  | SCI/Ctrl | 19 | 0.2602±0.01755 | SCI/Neurog2 vs. SCI/Ctrl | 0.0325 |
|  |  | SCI/Ngn2 | 13 | 0.5745±0.08110 |  |  |

Table S5

Multiple comparison for unpaired non-parametric one-way ANOVA with Kruskal-Walis test

Two-tailed Unpaired nonparametric t test

|  |  | **Group** | **Number** | **Mean intensity (Mean±SEM)** | **Multiple Comparisons** | ***P* value** |
| --- | --- | --- | --- | --- | --- | --- |
| Fig.  6.C. | MAP2 | Sham | 10 | 1238822±393890 | SCI/Neurog2 vs. Sham | 0.1987 |
|  |  | SCI/PBS | 9 | 742876±205162 | SCI/Neurog2 vs. SCI/PBS | 0.0244 |
|  |  | SCI/Ctrl | 11 | 192032±113007 | SCI/Neurog2 vs. SCI/Ctrl | <0.0001 |
|  |  | SCI/Ngn2 | 9 | 2076241±305695 |  |  |
| Fig.  6.D. | MAP2 in GFP | SCI/Ctrl | 9 | 20704±15908 | SCI/Neurog2 vs. SCI/Ctrl | 0.0003 |
|  |  | SCI/Ngn2 | 7 | 370709±106760 |  |  |
| Fig.  6.G. | GFAP | Sham | 10 | 332051±74824 | SCI/Neurog2 vs. Sham | 0.0575 |
|  |  | SCI/PBS | 7 | 1576571±171130 | SCI/Neurog2 vs. SCI/PBS | 0.0058 |
|  |  | SCI/Ctrl | 18 | 1320172±113411 | SCI/Neurog2 vs. SCI/Ctrl | 0.0035 |
|  |  | SCI/Ngn2 | 24 | 800887±7344 |  |  |
| Fig.  6.H. | NeuN | Sham | 4 | 364167±  92749 | SCI/Neurog2 vs. Sham | 0.9996 |
|  |  | SCI/PBS | 4 | 91905±  92749 | SCI/Neurog2 vs. SCI/PBS | 0.0245 |
|  |  | SCI/Ctrl | 13 | 77222±  92749 | SCI/Neurog2 vs. SCI/Ctrl | 0.0019 |
|  |  | SCI/Ngn2 | 6 | 356698±  92749 |  |  |
| Fig.  6.I. | NeuN in GFP | SCI/Ctrl | 66 | 297.5±  41.88 | SCI/Neurog2 vs. SCI/Ctrl |  |
|  |  | SCI/Ngn2 | 65 | 13466±  1842 |  |  |
| Fig.  6.J. | Isl1 | Sham | 14 | 385982±60547 | SCI/Neurog2 vs. Sham | 0.5639 |
|  |  | SCI/PBS | 4 | 42811±27259 | SCI/Neurog2 vs. SCI/PBS | 0.2820 |
|  |  | SCI/Ctrl | 14 | 91898±31917 | SCI/Neurog2 vs. SCI/Ctrl | 0.0317 |
|  |  | SCI/Ngn2 | 9 | 279578±92293 |  |  |
| Fig.  6.K. | Isl1 in GFP | SCI/Ctrl | 18 | 22888±11824 | SCI/Ngn2 vs. SCI/Ctrl | 0.0266 |
|  |  | SCI/Ngn2 | 12 | 46023±68353 |  |  |
| Fig.  6.L. | GFP in Isl1  (%) | SCI/Ctrl | 7 | 0±0 | SCI/Neurog2 vs. SCI/Ctrl | <0.0001 |
|  |  | SCI/Ngn2 | 6 | 44.07976±7.465214 |  |  |

Table S5

Multiple comparison for unpaired non-parametric one-way ANOVA with Kruskal-Walis test

| Fig.  7.C-E. | BrdU (3W) | **Group** | **Number** | **Mean±SEM** | **Multiple Comparisons** | ***P* value** |
| --- | --- | --- | --- | --- | --- | --- |
|  | GFAP+/BrdU+ (Ctrl) | GFP+ | 6 | 4.293±2.886 | GFP- vs. GFP+ | 0.0042 |
|  |  | GFP+GFAP+ | 6 | 4.293±2.886 | GFP- vs. GFP+GFAP+ | 0.0042 |
|  |  | GFP- | 6 | 95.71±2.886 |  |  |
|  | GFAP+/BrdU+ (Neurog2) | GFP+ | 5 | 28.48±18.89 | GFP- vs. GFP+ | 0.4511 |
|  |  | GFP+GFAP+ | 5 | 8.485±6.457 | GFP- vs. GFP+GFAP+ | 0.0947 |
|  |  | GFP- | 5 | 71.52±18.89 |  |  |
|  | NeuN+/BrdU+ (Ctrl) | GFP+ | 14 | 4.173±1.574 | GFP- vs. GFP+ | <0.0001 |
|  |  | GFP+GFAP+ | 14 | 0±0.000 | GFP- vs. GFP+GFAP+ | <0.0001 |
|  |  | GFP- | 14 | 95.83±1.574 |  |  |
|  | NeuN+/BrdU+ (Neurog2) | GFP+ | 11 | 4.385±1.120 | GFP- vs. GFP+ | 0.0006 |
|  |  | GFP+GFAP+ | 11 | 2.151±0.9320 | GFP- vs. GFP+GFAP+ | <0.0001 |
|  |  | GFP- | 11 | 95.62±1.120 |  |  |
|  | MBP+/BrdU+ (Ctrl) | GFP+ | 14 | 5.189±2.606 | GFP- vs. GFP+ | <0.0001 |
|  |  | GFP+GFAP+ | 14 | 0±0.000 | GFP- vs. GFP+GFAP+ | <0.001 |
|  |  | GFP- | 14 | 94.81±2.606 |  |  |
|  | MBP+/BrdU+ (Neurog2) | GFP+ | 13 | 6.447±4.460 | GFP- vs. GFP+ | <0.0001 |
|  |  | GFP+GFAP+ | 13 | 0±0.000 | GFP- vs. GFP+GFAP+ | <0.0001 |
|  |  | GFP- | 13 | 93.55±4.460 |  |  |

Table S6

Multiple comparison for unpaired non-parametric one-way ANOVA with Kruskal-Walis test

| Fig.  S5.  C-E. | BrdU (8W) | **Group** | **Number** | **Mean±SEM** | **Multiple Comparisons** | ***P* value** |
| --- | --- | --- | --- | --- | --- | --- |
|  | GFAP+/BrdU+ (Ctrl) | GFP+ | 7 | 12.06±5.997 | GFP- vs. GFP+ | 0.003 |
|  |  | GFP+GFAP+ | 7 | 9.683±4.258 | GFP- vs. GFP+GFAP+ | 0.002 |
|  |  | GFP- | 7 | 87.94±5.997 |  |  |
|  | GFAP+/BrdU+ (Neurog2) | GFP+ | 17 | 5.906±2.217 | GFP- vs. GFP+ | <0.0001 |
|  |  | GFP+GFAP+ | 17 | 1.186±0.8886 | GFP- vs. GFP+GFAP+ | <0.0001 |
|  |  | GFP- | 17 | 94.09±2.217 |  |  |
|  | NeuN+/BrdU+ (Ctrl) | GFP+ | 17 | 10.36±5.083 | GFP- vs. GFP+ | <0.0001 |
|  |  | GFP+GFAP+ | 17 | 0.3922±0.3922 | GFP- vs. GFP+GFAP+ | <0.0001 |
|  |  | GFP- | 17 | 89.64±5.083 |  |  |
|  | NeuN+/BrdU+ (Neurog2) | GFP+ | 23 | 9.82±4.517 | GFP- vs. GFP+ | <0.0001 |
|  |  | GFP+GFAP+ | 23 | 9.623±4.523 | GFP- vs. GFP+GFAP+ | <0.0001 |
|  |  | GFP- | 23 | 85.83±5.952 |  |  |
|  | MBP+/BrdU+ (Ctrl) | GFP+ | 6 | 10.34+3.729 | GFP- vs. GFP+ | >0.99 |
|  |  | GFP+GFAP+ | 6 | 0.000±0.000 | GFP- vs. GFP+GFAP+ | <0.001 |
|  |  | GFP- | 6 | 89.66±3.729 |  |  |
|  | MBP+/BrdU+ (Neurog2) | GFP+ | 13 | 12.01±3.684 | GFP- vs. GFP+ | 0.035 |
|  |  | GFP+GFAP+ | 13 | 0.000±0.000 | GFP- vs. GFP+GFAP+ | 0.000 |
|  |  | GFP- | 13 | 87.99±3.684 |  |  |
